# Reduced expression of the psychiatric risk gene DLG2 (PSD93) impairs hippocampal synaptic integration and plasticity

**DOI:** 10.1101/2021.08.02.454736

**Authors:** Simonas Griesius, Cian O’Donnell, Sophie Waldron, Kerrie L. Thomas, Dominic M. Dwyer, Lawrence S. Wilkinson, Jeremy Hall, Emma S. J. Robinson, Jack R. Mellor

## Abstract

**Background:** Genetic variations indicating loss of function in the *DLG2* gene have been associated with markedly increased risk for schizophrenia, autism spectrum disorder, and intellectual disability. *DLG2* encodes the postsynaptic scaffolding protein DLG2 (PSD93) that interacts with NMDA receptors, potassium channels, and cytoskeletal regulators but the net impact of these interactions on synaptic plasticity, likely underpinning cognitive impairments associated with these conditions, remains unclear. **Methods:** Hippocampal CA1 neuronal excitability and synaptic function were investigated in a novel clinically relevant heterozygous *Dlg2+/−* rat model using *ex vivo* patch-clamp electrophysiology, pharmacology, and computational modelling. **Results:** *Dlg2+/−* rats had increased NMDA receptor-mediated synaptic currents and, conversely, impaired associative long-term potentiation. This impairment resulted from an increase in potassium channel function leading to a decrease in input resistance and reduced supra-linear dendritic integration during induction of associative long-term potentiation. Enhancement of dendritic excitability by blockade of potassium channels or activation of muscarinic M1 receptors with selective allosteric agonist 77-LH-28-1 reduced the threshold for dendritic integration and 77-LH-28-1 rescued the associative long-term potentiation impairment in the *Dlg2+/−* rats. **Conclusions:** Despite increasing synaptic NMDA receptor currents, the combined impact of reduced DLG2 impairs synaptic integration in dendrites resulting in disrupted associative synaptic plasticity. This biological phenotype can be reversed by compound classes used clinically such as muscarinic M1 receptor agonists and is therefore a potential target for therapeutic intervention.

## Introduction

Genetic variations at the *DLG2* gene locus are linked to multiple psychiatric disorders including schizophrenia(1, 2), bipolar(3, 4), autism spectrum(5–7), attention deficit hyperactivity(8), intellectual disability(9, 10), and Parkinson’s disease(11, 12). This clinical evidence indicates the significance of DLG2 in the aetiology of psychopathologies common to a broad range of disorders and suggests core underlying mechanisms and biological phenotypes. Many of the genetic variations are predicted to produce a loss of function for DLG2 in one copy of the gene but the resulting changes in neuronal function are poorly understood(13–16).

DLG2 is a member of a family of membrane-associated guanylate kinase (MAGUK) proteins enriched at synaptic locations that encodes the scaffolding protein PSD93 (also referred to as DLG2 or Chapsyn-110). DLG2 interacts directly with a number of other proteins in the postsynaptic density of excitatory synapses, such as NMDA receptor (NMDAR) subunit GluN2B(17–20), AMPA receptor auxiliary subunit stargazin(21), potassium channels Kir2.3(22), Kir2.2(23) and Kv1.4(24), as well as proteins involved in potassium channel palmitoylation, cell adhesion, microtubule assembly, and cell signalling, palmitoyltransferase ZDHHC14(25), neuroligin1-3(17), Fyn(26, 27), ERK2(28), GKAP(29), and MAP1A(30). Uniquely to the MAGUK family, DLG2 is targeted to the axon initial segment where it regulates neuronal excitability via its interactions with potassium channels(25, 31). At a functional level, homozygous *Dlg2−/−* knockout mice have altered glutamatergic synapse function(32–35) and impaired long-term potentiation (LTP) in the hippocampus(33). These synaptic perturbations could underlie the common cognitive psychopathologies of the psychiatric disorders associated with *DLG2*. Indeed, *Dlg2−/−* mice have been shown to have impaired performance in the object-location paired associates learning task(36). Homozygous *Dlg2−/−* mice also exhibit increased grooming behaviour(35) and altered social interaction but without consistent effects in negative valence tasks such as the open field test(35, 37).

Impaired synaptic plasticity resulting from the loss of DLG2 is a potential biological phenotype underpinning trans-diagnostic cognitive psychopathologies but the mechanism by which reduced DLG2 expression leads to impaired synaptic plasticity is unclear. Furthermore, mechanistic understanding for the impact of DLG2 loss may reveal new targets for therapeutic intervention. Most animal models for DLG2 loss have employed full knockouts of the gene but these do not accurately represent the heterozygous nature of DLG2 genetic variants in patient populations and potentially engage compensatory expression by other MAGUK proteins(32) (38) that is not present in heterozygous reduced gene dosage models (Fig 1A-C). Therefore, here we investigate the combined impact of low gene dosage DLG2 on synaptic function, neuronal excitability and morphology using a novel CRISPR-Cas9 engineered heterozygous *Dlg2+/−* (het) rat model to understand the interactions that lead to impaired synaptic plasticity and cognitive function.

**Figure 1.**
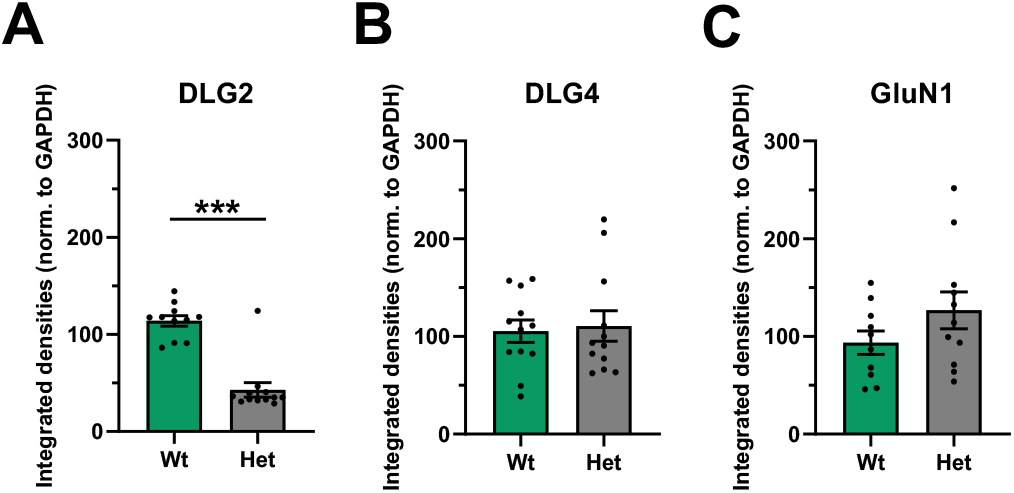
DLG2 expression is reduced in *Dlg2+/−* hets, with no change in DLG4 or GluN1 expression. Expression of DLG2 (unpaired t-test: P < 0.001) (A), DLG4 (unpaired t-test: P = 0.784) (B), and GluN1 (unpaired t-test: P = 0.163) (C) in the hippocampus of *Dlg2+/−* het and wt rats. Data from 23 (DLG2), 24 (DLG4), and 21 (GluN1) rats. Summary values depicted as mean ± SEM. * P < 0.05, ** P < 0.01, *** P < 0.001 (unpaired t-test)

## Methods and materials

### Animals and husbandry

All procedures were carried out under local institutional guidelines, approved by the University of Bristol Animal Welfare and Ethical Review Board, and in accordance with the UK Animals (Scientific procedures) Act 1986. The experiments employed a novel *Dlg2+/−* heterozygous rat model generated on a Hooded Long Evans background using CRISPR-Cas9 genomic engineering that targeted a 7bp deletion to exon 5 of the rat *Dlg2* gene, resulting in a downstream frame shift in exon 6 and the production of a premature stop codon that led to a reduction in Dlg2 protein levels in the hippocampus (Fig 1). Full details of the creation, quality control and off-target assessment of the *Dlg2+/−* model can be found in Supplement 1. Male *Dlg2+/−* rats were bred with wild type (wt) female rats, generating mixed litters of *Dlg2+/−* and wt littermate offspring. The *Dlg2+/−* animals were viable and showed no signs of ill health, with normal litter sizes containing the expected Mendelian ratio of positive to wt genotypes and normal sex ratios, there were no effects in survival of the *Dlg2+/−* rats to adulthood and no effects on general morbidity, including fertility, or mortality throughout the lifespan. Further details of animal husbandry, breeding strategy and viability are described in Supplement 1. Approximately equal numbers of each sex rats aged P50-75 were used, with experimenter blind to genotype during experiments and data analysis.

### Brain slice preparation, electrophysiology, protein quantification and computational modelling

These procedures are described in Supplement 1.

### Statistical analysis

3-way and 2-way ANOVA, 3-way repeated measures ANOVA, Komolgorov-Smirnov test, as well as paired and unpaired t-tests were used as appropriate, with full statistical results available in Supplement 2. Genotype, sex, and dorsal-ventral aspects of the hippocampus, as well as repeated measurements, were factored into all analyses, as appropriate. Genotype was viewed as the primary output factor shown in the figures but where effects of other factors were found, the data are presented in Supplement 1 (Figs S6, S10, S12, S13, S15, S16). Inclusion of animals in the analysis of a subset of experiments using multi-level general linear mixed modelling did not affect the statistical results indicating that the major source of variability arose between cells rather than animals. Therefore, cell was defined as the experimental unit and we report numbers of cells and animals in figure legends. α = 0.05 was applied for all tests, except the Kolmogorov-Smirnov test where α = 0.01 was applied. The degrees of freedom, F, and P values are presented in the text, figures, and Supplement 2.

## Results

*Dlg2+/−* heterozygous knockout rats were generated by CRISP-Cas9 targeting of the *Dlg2* gene (Supplement 1). In this model, DLG2 protein expression levels were reduced by ∼50% in hippocampus without effects on expression of other components of the postsynaptic density, including the closely related MAGUK DLG4 (PSD95) and the GluN1 NMDAR subunit (Fig 1A-C).

### NMDAR currents

Protein-protein interaction studies have reported DLG2 to interact directly with NMDAR subunits(17–19) and with AMPAR indirectly(21). Further, DLG2 has also been shown to affect glutamatergic function in homozygous *Dlg2−/−* models, albeit with variable results in AMPA/NMDA ratio(32–34). To investigate whether glutamatergic function was affected in *Dlg2+/−* rats, the AMPA/NMDA ratio was measured in the CA1 region of hippocampal slices (Fig 2A). *Dlg2+/−* hets had a reduced AMPA/NMDA ratio in the SC pathway, with no effect in the TA pathway (Fig 2B-D). AMPAR-mediated miniature excitatory postsynaptic currents (mEPSCs, Supplement 1 Fig S3) resulting from the activity of single synapses were recorded to probe which component of the AMPA/NMDA ratio was affected. mEPSCs from synapses on apical dendrites are more detectable than those from more distal synapses due to signal attenuation(39) so mEPSCs are assumed to arise from synapses in the SC pathway. There was no difference in the distributions of mEPSC amplitude, interevent interval, or decay tau across genotype (Fig 2E-I). Paired-pulse facilitation, measured in the AMPA/NMDA ratio experiment, was also not different across genotype in either pathway (Supplement 1 Fig S5). Together, these results show no change in postsynaptic AMPAR function and presynaptic glutamate release probability in the SC pathway. It follows that the AMPA/NMDA ratio effect in the SC pathway was due to an increase in NMDAR function. This could result from either an increase in NMDAR number or a change in subunit composition between GluN2A and GluN2B. To test subunit composition, NMDAR currents were isolated (Fig 2J) and the selective GluN2B negative allosteric modulator RO256981 was applied. RO256981 decreased EPSC amplitude (Fig 2J-L) and increased the EPSC decay time in both the SC and the TA pathways (Fig 2M-N). There was a trend toward a genotype x drug interaction in the EPSC amplitude measurement in the SC pathway but no genotype x drug interaction in the EPSC decay kinetics. Together, these results show similar NMDAR subunit composition across genotype and therefore the enhancement in NMDAR function likely arises from increased receptor numbers at SC synapses.

**Figure 2.**
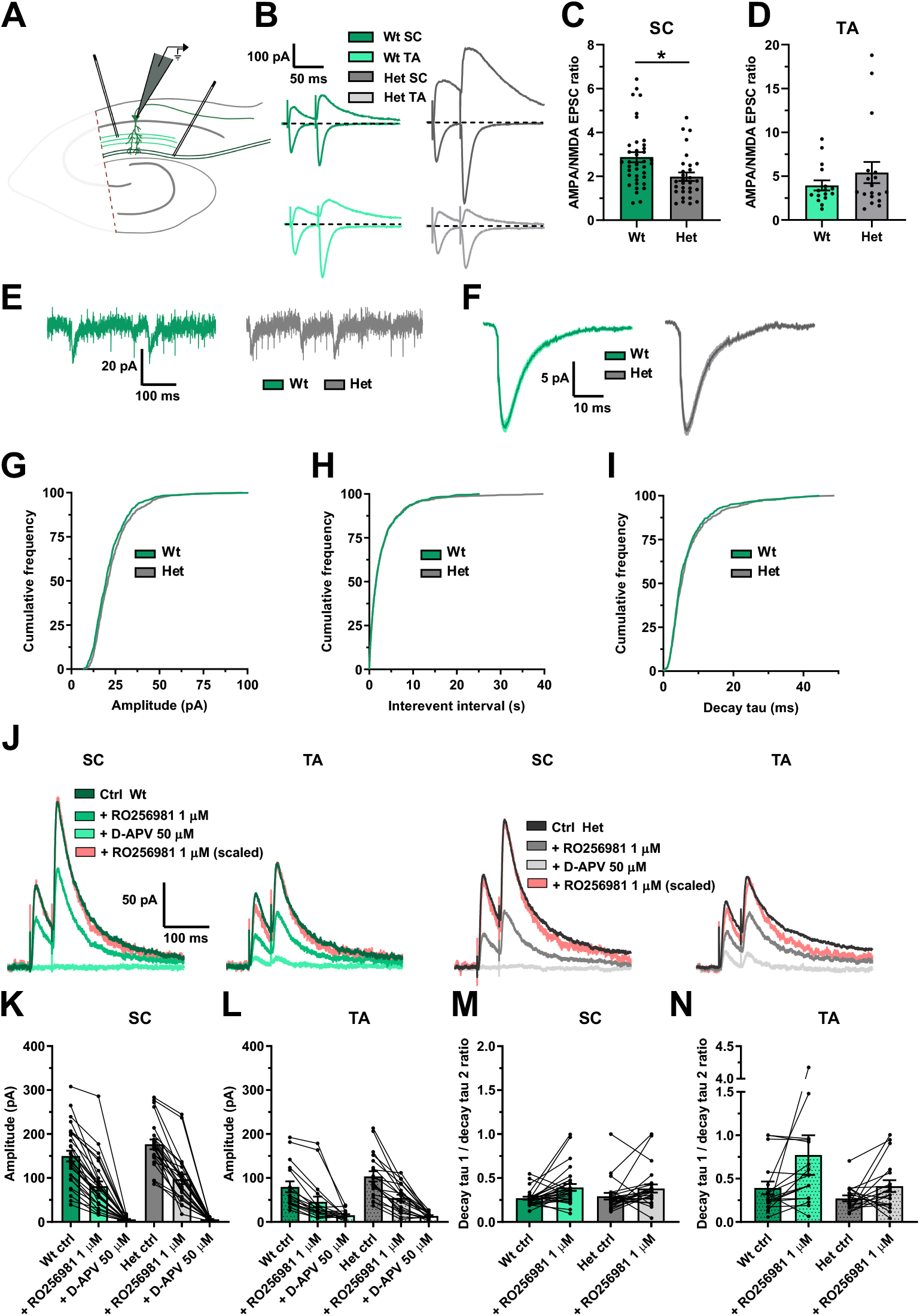
Glutamatergic function is altered in the *Dlg2+/−* hets due to an increase in NMDAR current, with no change in AMPAR current or GluN2b subunit expression. **A)** Schematic representation of the hippocampal slice recording setup, with the CA3 removed and stimulating electrodes in the stratum radiatum and in the stratum lacunosum moleculare. **B)** AMPA/NMDA EPSC ratio example traces. The primarily AMPAR-mediated EPSCs were recorded at a holding potential of -70 mV, whilst the AMPA-and NMDA-mediated EPSCs were recorded at a holding potential of 40 mV. The AMPA/NMDA ratio was derived by dividing the peak EPSC amplitude at -70 mV by the EPSC amplitude 45 ms after stimulation at 40 mV. AMPA/NMDA EPSC ratio across genotype in the SC (3-way ANOVA: genotype main effect: F _1, 68_ = 4.791, P = 0.033) **(C)** and the TA (3-way ANOVA: genotype main effect: F _1, 34_ = 0.583, P = 0.452) **(D)** pathways. 68 cells from 32 animals for the SC data set and 34 cells from 16 animals for the TA data set. Example mEPSC traces **(E)** and mean mEPSCs **(F)** across genotype. Cumulative frequency plots of amplitude (Kolmogorov–Smirnov Test: P = 0.028) **(G)**, interevent interval (Kolmogorov–Smirnov Test: P = 0.999) **(H)**, and decay tau (Kolmogorov–Smirnov Test: P = 0.067) **(I)**. 45 cells from 14 animals. **J)** GluN2b example EPSC traces. The traces following RO256981 1µM administration were also peak scaled to better illustrate changes in decay kinetics. EPSC amplitude in the SC (3-way repeated-measures ANOVA: drug main effect: F _2, 88_ = 260.603, P < 0.001. Genotype x drug interaction: F _2, 88_ = 2.952, P = 0.057) **(K)** and TA (3-way repeated-measures ANOVA: drug main effect: F _2, 56_ = 42.076, P < 0.001. Genotype x drug interaction: F _2, 56_ = 1.738, P = 0.185) **(L)** pathways. Decay tau 1 / decay tau 2 ratio in SC (3-way repeated-measures ANOVA: drug main effect: F _1, 45_ = 9.715, P = 0.003. Genotype x drug interaction: F _1, 45_ = 0.272, P = 0.605) **(M)** and TA (3-way repeated-measures ANOVA: drug main effect: F _1, 27_ = 3.410, P = 0.076. Genotype x drug interaction: F _1, 27_ = 0.831, P = 0.370) **(N)** pathways. 53 cells from 14 animals (SC), 38 cells from animals 14 (TA). Summary values depicted as mean ± SEM. * P < 0.05, ** P < 0.01, *** P < 0.001 (3-way ANOVA between subject effect)

### Synaptic integration and aLTP

NMDARs contribute to dendritic integration of spatiotemporally coherent inputs that can summate supralinearily to drive associative LTP (aLTP) relevant to the learning of novel representations(40–47). Given increased NMDAR function, it is predicted that dendritic integration and plasticity would be facilitated in the *Dlg2+/−* hets. To test this, aLTP was induced by stimulating SC and TA pathways simultaneously (Fig 3A-B). An additional independent SC pathway was also stimulated as a negative control and a pathway check was done to confirm pathway independence (Supplement 1 Fig S4). Despite the increased NMDAR function in the *Dlg2+/−* hets, the induction protocol resulted in reduced aLTP in the *Dlg2+/−* hets in both SC and TA test pathways but robust aLTP in the wts (Fig 3C-E). Critically, baseline EPSC amplitudes were not different (Fig 3G), suggesting that all neurons received similar inputs. However, during induction (Fig 3F) the number of elicited action potential bursts and single spikes was reduced in the *Dlg2+/−* hets (Fig 3H,J) and there was a trend suggesting reduced overall depolarisation in response to synaptic stimulation (Fig 3I). Both spike number and depolarisation during induction correlated with LTP in the SC pathway but not the TA pathway (Supplement 1 Fig S7). There was no effect of genotype on the after-hyperpolarisation (Fig 3K). This suggests that although synaptic receptor numbers are increased in the *Dlg2+/−* hets, the integration of synaptic inputs from the SC and TA pathways is impaired, reducing dendritic depolarisation and action potential spiking which are the drivers of aLTP.

**Figure 3.**
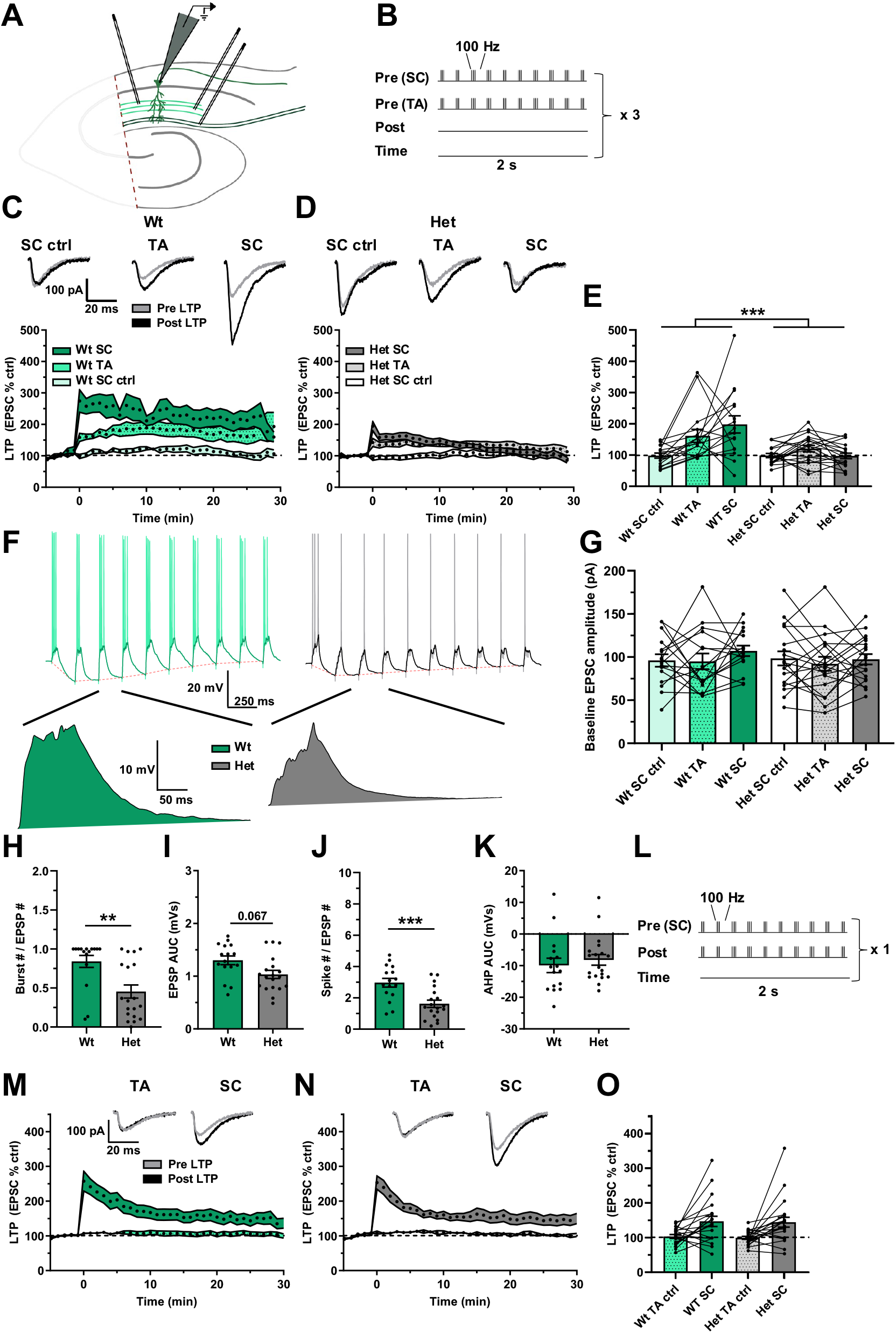
Attenuated aLTP in the *Dlg2+/−* hets despite normal TBS LTP. **A)** Schematic representation of the hippocampal slice recording setup, with the CA3 removed and stimulating electrodes in two separate areas of the stratum radiatum and in the stratum lacunosum moleculare. **B)** aLTP induction protocol, where one SC pathway and one TA pathway were tested and where the second SC pathway acted as a negative control. There was no induced somatic depolarisation. This induction protocol was repeated thrice at an interval of 10 seconds. aLTP over time in wts **(C)** and *Dlg2+/−* hets **(D)**. Example traces pre- and post-induction are displayed for the wt and het groups above their corresponding plots of LTP over time. **E)** aLTP at the 25-30 minute mark post induction across genotype and pathway (3-way repeated-measures ANOVA: pathway effect: F _2, 54_ = 7.300, P = 0.002. Genotype main effect: F _1, 27_ = 15.687, P < 0.001. Genotype x pathway interaction: F _2, 54_ = 7.376, P = 0.001). **F)** Example traces of LTP induction, with example EPSPs following *post hoc* spike truncation. **G)** Baseline EPSC amplitude across genotype (3-way repeated-measures ANOVA: pathway effect: F _2, 54_ = 1.227, P = 0.301. Genotype main effect: F _1, 27_ = 0.553, P = 0.463. Genotype x pathway interaction: F _2, 54_ = 0.344, P = 0.711). Burst number (3-way ANOVA: genotype main effect: F _1, 27_ = 10.407, P = 0.003) **(H)**, EPSP AUC (3-way ANOVA: genotype main effect: F _1,27_ = 3.63, P = 0.067) **(I),** total spike number (3-way ANOVA: genotype main effect: F _1, 27_ = 15.877, P < 0.001) **(J)**, and afterhyperpolarisation AUC (3-way ANOVA: genotype main effect: F _1, 27_ = 0.036, P = 0.85) **(K)** across genotype during LTP induction. 35 cells from 17 animals. **L)** Theta burst LTP induction protocol, where the SC pathway was paired with somatic depolarisation and where the TA pathway acted as a negative control. Theta burst LTP over time in wts **(M)** and *Dlg2+/−* hets **(N)**. **O)** Theta burst LTP at the 25-30 minute mark post induction across genotype (3-way repeated-measures ANOVA: pathway effect: F _1, 33_ = 18.979, P < 0.001. Genotype main effect: F _1, 33_ = 0.04, P = 0.843. Genotype x pathway interaction: F _1, 33_ = 0.004, P = 0.950). 41 cells from 19 animals. Summary values depicted as mean ± SEM. * P < 0.05, ** P < 0.01, *** P < 0.001 (3-way ANOVA between subject effect)

To test the necessity of action potentials for aLTP, a paired theta burst LTP induction protocol was used, where action potentials were driven by somatic current injection to bypass dendritic integration, and spikes were paired with simultaneous SC pathway stimulation (Fig 3L). Under these conditions, robust LTP was induced in the SC pathway in both genotypes, with the TA pathway acting as negative control (Fig 3M-O). Similarly, when aLTP was tested using baseline EPCSs doubled in amplitude, maximal LTP was induced and there was no effect of genotype (Supplement 1 Figs S8&S9). This indicates that the hets are fundamentally able to undergo LTP but their ability to integrate inputs is impaired.

To directly test synaptic integration, the number of activated synapses required to generate supra-linear summation of EPSPs was assessed, which is a measure of the ability for synapses to integrate within dendrites(44, 45, 47). To activate increasing numbers of synapses, the SC pathway was stimulated with increasing intensity. The number of activated synapses was measured by the slope of a single EPSP and the integration of synapses assessed by the amplitude and duration of a compound summated EPSP (area under the curve – AUC) elicited by repetitive high frequency synaptic stimulation (Fig 4A). As stimulation intensity was increased the number of active synapses increased in a linear relationship with the amplitude and durations of the summated compound EPSP until a “change point” was reached (see methods) after which the relationship became supra-linear because the duration of the compound EPSP increased (Fig 4B), indicative of activation of regenerative or plateau potentials within the dendrites(44, 45, 47). The inhibition of these regenerative potentials by D-APV demonstrated their dependence on NMDAR activation (Fig 4A-B). The change point was increased in the *Dlg2+/−* hets (Fig 4B-C), indicating that het neurons required more synaptic inputs to undergo the transition to supra-linear integration. Additionally, the maximum duration of the compound EPSP as a ratio to the corresponding slope of the single EPSP was reduced in the *Dlg2+/−* hets (Fig 4D). This again indicates that *Dlg2+/−* hets require more synaptic input to integrate dendritic inputs and produce the supra-linear regenerative potentials important for aLTP.

**Figure 4.**
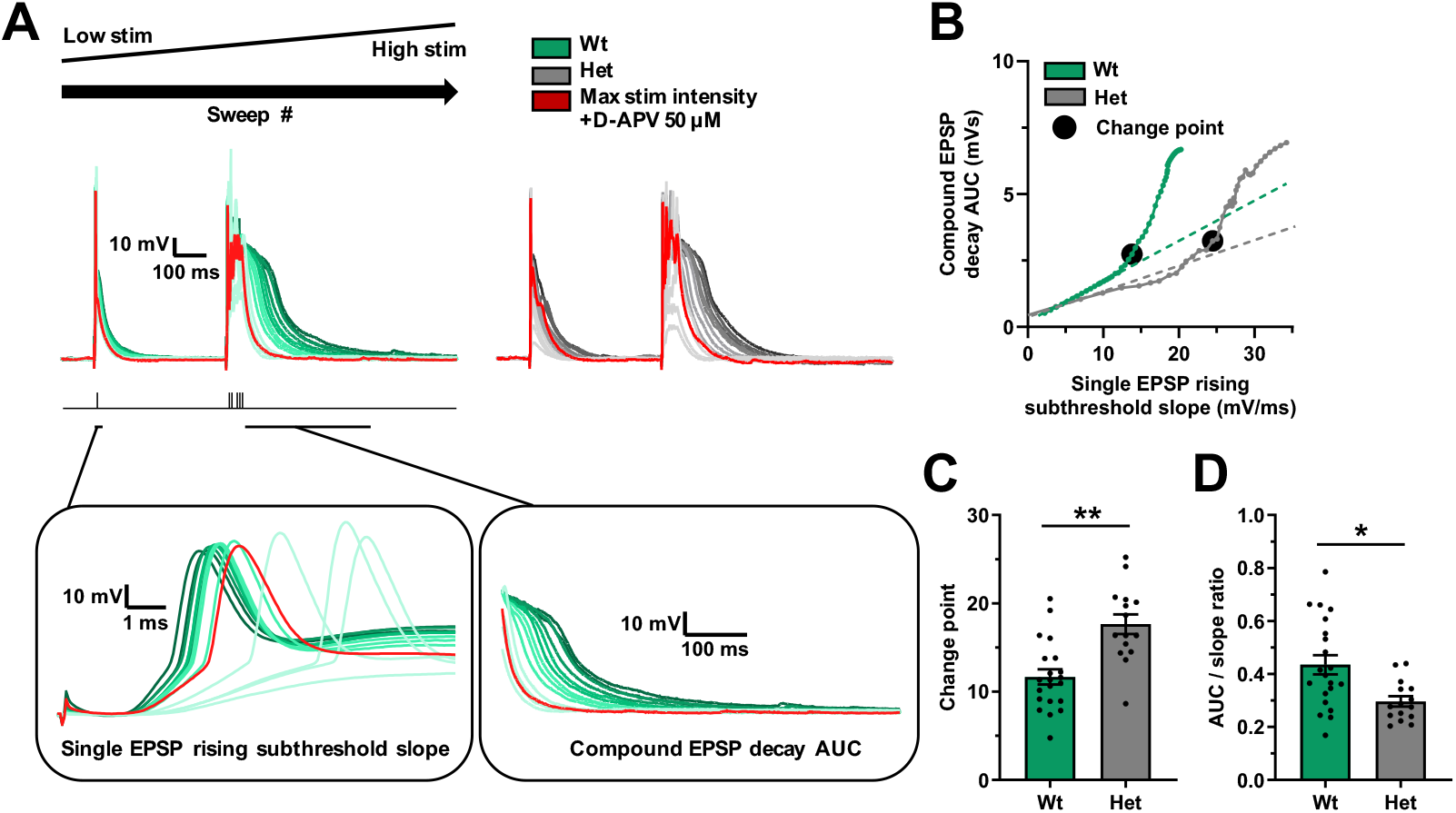
Increased threshold for supra-linear dendritic integration in the *Dlg2+/−* hets in the SC pathway. **A)** Example traces depicting a single EPSP followed by a compound EPSP at increasing stimulation intensities (light to dark) over consecutive recording sweeps. The red trace represents the response at maximal stimulation intensity in the presence of 50 µM D-APV. The left inset depicts a zoomed-in view of the single EPSP, where rising subthreshold slope was measured. The right inset depicts a zoomed-in view of the compound EPSP decay, where the AUC was measured. **B)** Example relationships between the single EPSP rising subthreshold slope and the compound EPSP decay AUC across genotype, with the change points marked by black circles. Change point (EPSP rising subthreshold slope mV/mS) (3-way ANOVA: genotype main effect: F _1, 34_ = 12.625, P = 0.001) **C)** and AUC/slope ratio (3-way ANOVA: genotype main effect: F _1, 34_ = 8.003, P = 0.009) **(D)** across genotype. 38 cells from 23 animals. Summary values depicted as mean ± SEM. * P < 0.05, ** P < 0.01, *** P < 0.001 (3-way ANOVA between subject effect)

### Mechanism for impaired synaptic integration and plasticity

Reduced synaptic integration in dendrites could arise from multiple mechanisms. Based on previous findings in CA1 pyramidal neurons the three most likely are: i) Reduced expression of hyperpolarisation-activated cyclic nucleotide-gated (HCN) channels that regulate neuronal excitability and contribute to dendritic integration(48–50), ii) Increased expression of small conductance calcium-activated potassium (SK) channels that inhibit NMDARs at synapses, reducing dendritic integration and LTP(51–53), iii) Reduced input resistance by increased potassium channel expression particularly in dendritic locations to reduce dendritic integration and LTP(44, 45, 54–58). Although blocking HCN channels produced robust effects on neuronal excitability (including spiking, sag, and input resistance) there were no differential effects across genotype (Fig 5A-F). There were also no genotype-specific effects on cellular resonance or impedance that are directly dependent on HCN channels(59–63) (Supplement 1 Fig S11). As previously described, the SK channel blocker apamin produced an increase in EPSP duration in the SC and TA pathways (Fig 5G-I) indicating increased NMDAR activation during synaptic stimulation(51–53). However, the regulation of synaptic NMDAR function by SK channels was similar between genotypes indicating no change in SK channel expression. Therefore, differential HCN or SK channel function is unlikely to explain the difference in synaptic integration between genotypes.

**Figure 5.**
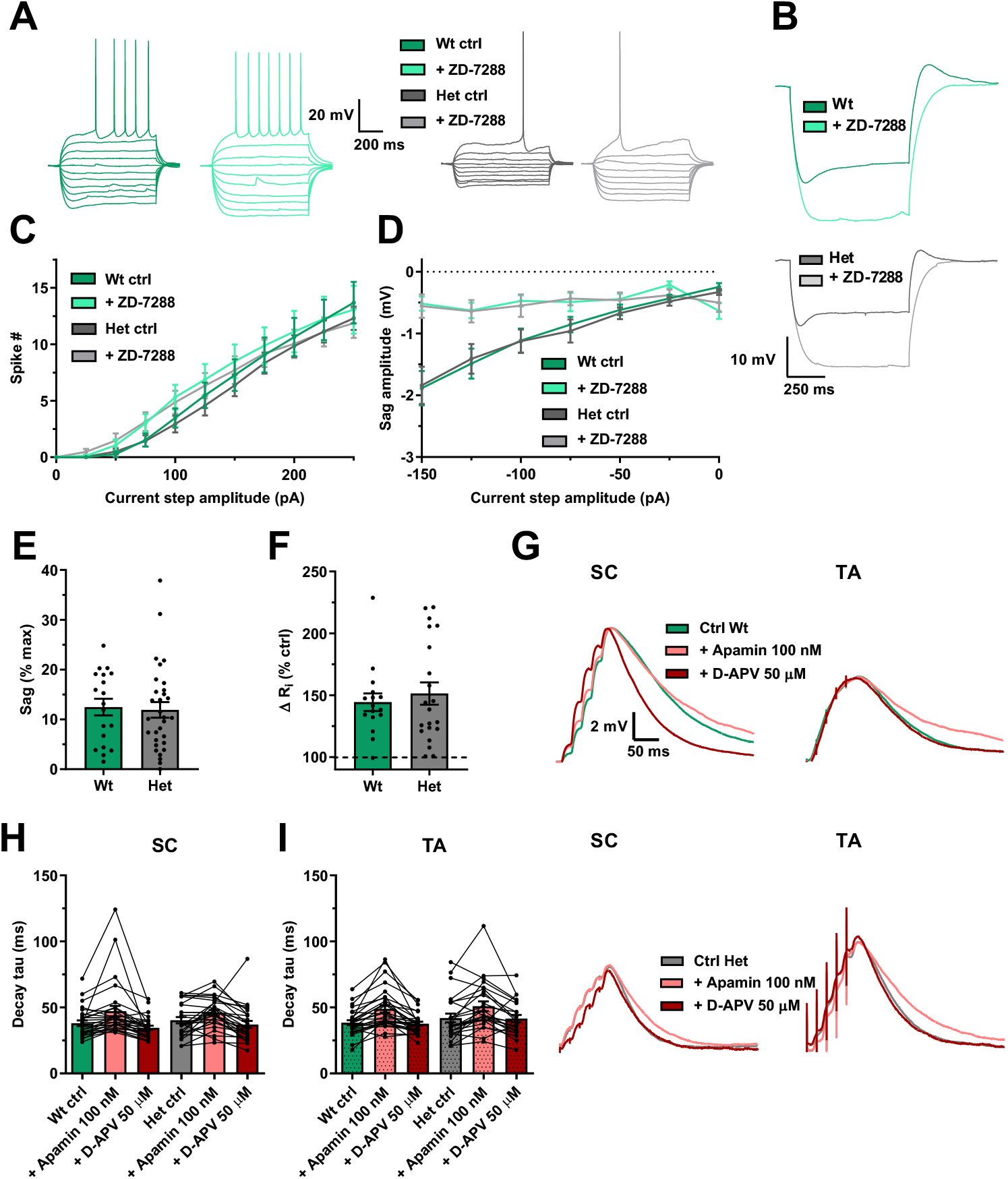
Unaltered HCN and SK channel function in the *Dlg2+/−* hets. **A)** Example voltage traces in response to current steps (-150 to 250 pA) before and after application of ZD-7288 20 µM. **B)** Voltage traces in response to a hyperpolarising current step of -150 pA before and after the application of ZD-7288 20 µM. Spike number (3-way repeated-measures ANOVA: drug effect: F _1, 28_ = 0.321, P = 0.576. Current step effect: F _10, 280_ = 99.423, P < 0.001. Genotype main effect: F _1, 28_ = 0.272, P = 0.606. Drug x genotype interaction: F _1, 28_ = 0.304, P = 0.586. Drug x step interaction: F _10, 280_ = 4.126, P < 0.001. Drug x step x genotype interaction: F _10, 280_ = 0.134, P = 0.999) **(C)** and sag amplitude (3-way repeated-measures ANOVA: drug effect: F _1, 28_ = 14.797, P = 0.001. Current step effect: F _6, 168_ = 14.072, P < 0.001. Genotype main effect: F _1, 28_ = 0.03, P = 0.864. Drug x genotype interaction: F _1, 28_ = 0.206, P = 0.654. Drug x step interaction: F _10, 280_ = 4.126, P < 0.001. Drug x step x genotype interaction: F _10, 280_ = 0.564, P = 0.758) **(D)** across genotype and before and after the application of ZD-7288 20 µM. Sag as a percentage of max voltage deflection (3-way repeated-measures ANOVA: genotype main effect: F _1, 50_ = 0.182, P = 0.672) **(E)** and change in input resistance (3-way repeated-measures ANOVA: genotype main effect: F _1, 37_ = 0.351, P = 0.558) **(F)** following the application of ZD-7288 20 µM. 37 cells from 18 animals. **G)** EPSPs before and after the application of apamin 100 nM. EPSP decay tau across genotype in SC (3-way repeated-measures ANOVA: drug effect: F _2, 92_ = 18.327, P < 0.001. Genotype main effect: F _1, 46_ = 0.057, P = 0.812. Drug x genotype interaction: F _2, 92_ = 0.76, P = 0.471.) **(H)**, and TA (3-way repeated-measures ANOVA: drug effect: F _2, 86_ = 29.346, P < 0.001. Genotype main effect: F _1, 43_ = 0.391, P = 0.535. Genotype x drug interaction: F _2, 86_ = 0.166, P = 0.847). **(I)** pathways. 55 cells from 17 animals (SC), 54 cells from animals 17 (TA). Summary values depicted as mean ± SEM. * P < 0.05, ** P < 0.01, *** P < 0.001 (3-way ANOVA between subject effect)

To assess input resistance, measurements were analysed from voltage clamp experiments (using identical conditions to the LTP experiments in Fig 3) and in current clamp experiments. In both these data sets *Dlg2+/−* hets had reduced input resistance (Fig 6A-D). This increase in electrical leak in the *Dlg2+/−* hets is predicted to reduce cross-talk between synapses and their integration leading to a reduced spike output but it is also expected to reduce the spike output in response to somatic current injection. However, despite reduced input resistance in the *Dlg2+/−* hets, there was no effect of genotype on spike output to current injection (rheobase) (Fig 6E-F). This could be explained by a depolarised resting membrane potential (Fig 6G) and a trend towards hyperpolarised action potential spike threshold (Fig 6H) in the *Dlg2+/−* hets indicating that smaller membrane potential depolarisations were required to initiate spikes. There was no effect of genotype on spike half-width, maximum spike slope, spike amplitude, or capacitance, and a slight decrease in latency to spike in the *Dlg2+/−* hets (Supplement 1 Fig S14).

**Figure 6.**
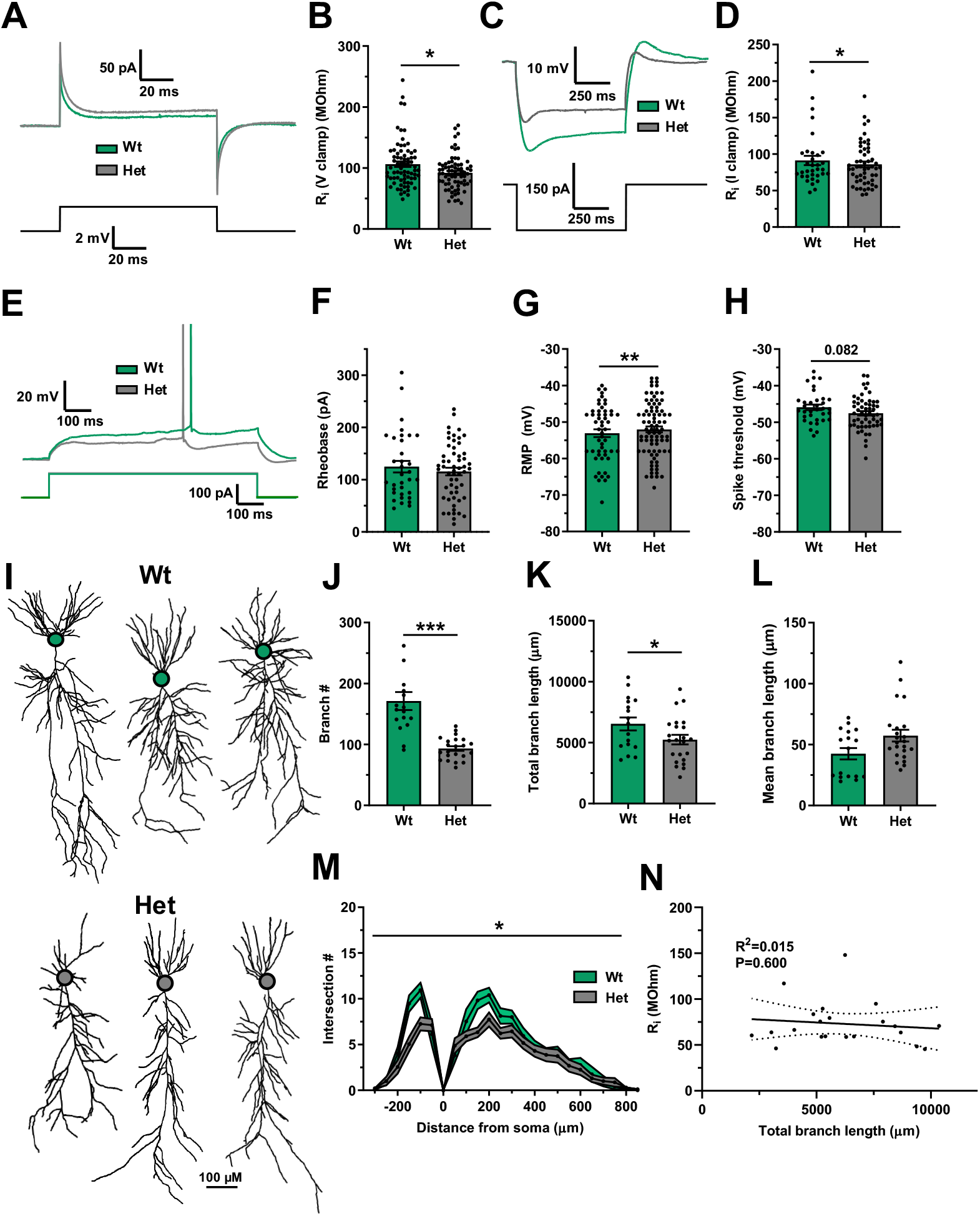
Reduced input resistance and dendritic arborisation in the *Dlg2+/−* hets. **A)** Current traces in response to a 2 mV voltage step for wt and het. **B)** Input resistance (V clamp) across genotype (3-way ANOVA: genotype main effect: F _1, 146_ = 5.698, P = 0.018). 146 cells from 58 animals. **C)** Voltage traces in response to a -150 pA current step across genotype. **D)** Input resistance (I clamp) across genotype (3-way ANOVA: genotype main effect: F _1, 78_ = 4.209, P = 0.044). **E)** Voltage deflections in response to a current step of equal size across genotype. Rheobase (3-way ANOVA: genotype main effect: F _1, 90_ = 0.011, P = 0.916) **(F)**, resting membrane potential (RMP) (3-way ANOVA: genotype main effect: F _1, 136_ = 7.075, P = 0.009) **(G)**, and spike threshold (3-way ANOVA: genotype main effect: F _1, 89_ = 3.105, P = 0.082) **(H)** across genotype. ∼136 cells from 38 animals. **(I)** Example morphological reconstructions across genotype, cell bodies denoted with circles. Dendritic branch number (3-way ANOVA: genotype main effect: F 1, 40 = 23.279, P < 0.001) **(J)**, total dendritic branch length (3-way ANOVA: genotype main effect: F 1, 40 = 7.002, P = 0.013) **(K)**, and mean dendritic branch length (3-way ANOVA: genotype main effect: F 1, 40 = 0.133, P = 0.718) **(L)** across genotype. **M)** Scholl analysis across genotype (3-way ANOVA: genotype main effect: F 1, 31 = 5.532, P = 0.025). 40 cells from 20 animals. **N)** Correlation between total dendritic branch length and input resistance dataset (Pearson correlation: R^2^ = 0.015, P = 0.600). Summary values depicted as mean ± SEM. * P < 0.05, ** P < 0.01, *** P < 0.001 (3-way ANOVA between subject effect)

Reduced input resistance in the *Dlg2+/−* hets could be explained via two mechanisms: i) increased membrane area through greater dendritic branching and extent(45, 64) or ii) increased membrane conductance, most likely caused by increased potassium channel expression. To test the first mechanism, a subset of neurons from the intrinsic excitability experiments were filled with neurobiotin to allow *post hoc* morphological analysis. Analysis of these neurons revealed that het neurons were smaller than wt neurons (Fig 6I) and had reduced dendritic branch number and total dendritic branch length but had similar mean dendritic branch lengths (Fig 6J-L). Scholl analysis demonstrated that *Dlg2+/−* het neurons had reduced dendritic arborisation overall, with the most striking differences in the basal and proximal apical regions (Fig 6M). Contrary to the predicted neuronal size – input resistance relationship, there was no correlation between total dendritic branch length and input resistance (Fig 6N). Therefore, reduced neuronal arborisation in the *Dlg2+/−* hets cannot explain the observed reduced input resistance and instead increased potassium channel expression is the most likely explanation.

To explore which potassium channels were most likely to reduce input resistance and enhance synaptic integration, a computational approach was employed where 6 representative reconstructed pyramidal neurons (3 wt, 3 het) were populated with voltage dependent sodium (NaV) and potassium (Ka, Kir, Kdr, and Km, A-type, inward rectifier, delayed rectifier and M-type respectively) channels as well as a voltage independent leak conductance. Computer simulations of these het reconstructed neurons showed consistently increased input resistance relative to wt (Fig 7A-B). This result was expected from the reduced dendritic arborisation and supports the conclusion that the *Dlg2+/−* hets have increased ionic conductances. DLG2 interacts with potassium inward rectifier Kir2.3(22) and Kir2.2(23) as well as A-type Kv1.4(24) channels which therefore represented the most likely channels to underpin decreased input resistance and increased synaptic integration. In order to probe the effects of these channels on input resistance, the wt reconstruction was populated with either Kir or Ka channels and channel conductances were scaled systematically over a ten-fold range (Fig 7C). The inclusion of Ka channels reduced input resistance with greater channel density causing greater reductions in input resistance. In contrast, the inclusion of Kir channels had no effect across the conductance ranges explored (Fig 7C). These potassium channel selective effects on input resistance map onto the voltage-dependent activation curves of these channels (Supplement 1 Fig S17). Based on these results, an increase in Ka channel expression or function is a good candidate to underly the decrease in input resistance in the *Dlg2+/−* hets. Next, to test the effects of input resistance, as well as Ka and Kir channels, on dendritic integration, a series of simulations were executed (Fig 7D). Dendrites were randomly activated in a cumulative manner, with increasing numbers of proximal apical and tuft dendrites being activated at any one time. Reminiscent of the dendritic integration experiment in slices (Fig 4), change points were observed where the relation between activated dendrite number and summated EPSP amplitude and duration transitioned from linear to supra-linear (Fig 7E). This change point shifted leftward with increasing numbers of activated tuft dendrites, indicating the facilitation of dendritic integration in the presence of more synaptic inputs. Similar to slice experiments, simulation of proximal apical dendrite activation alone was also able to generate supra-linear summation (Fig 7E). Following the inclusion of Ka channels into the simulation, supra-linear summation was abolished (Fig 7F) whereas the inclusion of Kir channels had no effect (Fig 7G) again reflecting the respective voltage-dependent activation curves for Ka and Kir channels (Supplement 1 Fig S17). Doubling the leak conductance, which approximately halved input resistance, considerably attenuated EPSP summation and shifted the change point to the right indicating reduced synaptic integration and supra-linearity (Fig 7H). Of the potassium channels interacting with DLG2, these results suggest that A-type potassium channels are the most likely candidates upregulated in the *Dlg2+/−* hets to underly the dendritic integration deficits.

**Figure 7.**
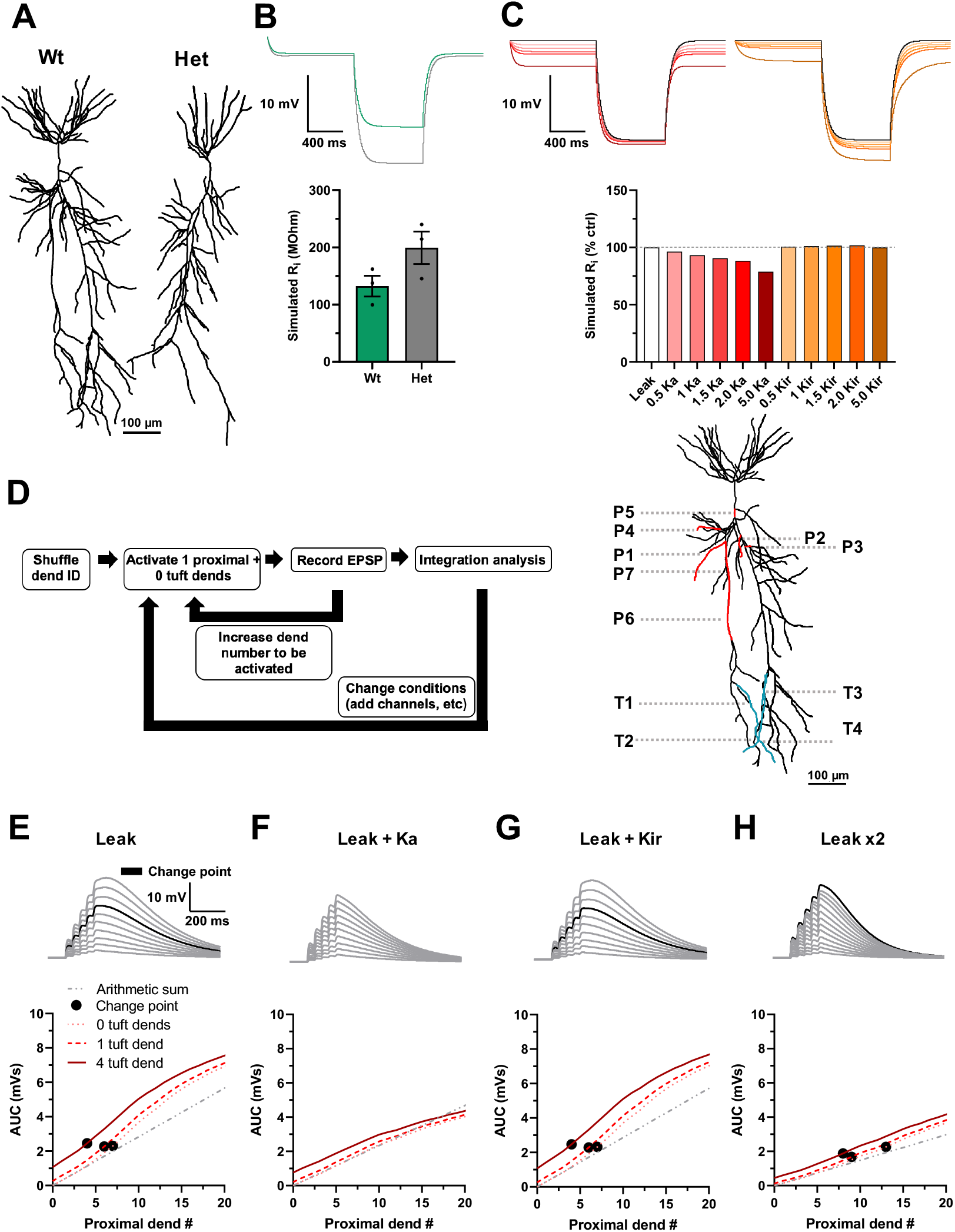
With equal ionic conductance, simulated het neurons have higher input resistance. Ka channels reduce input resistance and inhibit dendritic integration. **A)** Example reconstructions used in simulations. **B)** Simulated voltage traces in response to a -150 pA current step across genotype (top) and the corresponding input resistance (bottom). 6 neuronal reconstructions were populated with Ka, Kir, Km, Kdr, NaV channels. **C)** Voltage traces in response to a -150 pA current step in a wt reconstruction populated either with the Ka (pink) or the Kir (orange) channels (top). Channel conductance was scaled by a factor of 0.5-5 for each condition. Corresponding input resistance (bottom). **D)** Overview of the dendritic integration simulations. Different proportions of proximal and tuft dendrites were activated in a random, but consistent across experiments, sequence with 1 synapse per dendrite. AUC was calculated to identify the change point to supralinearity, conditions were altered, and the procedure was repeated. The first 7 proximal (P) (red) and first 4 tuft (T) (blue) activated dendrites are shown on top of the wt reconstruction used in the dendritic integration simulations, with the numbered dendrites indicating their relative activation sequence. EPSP AUC as a function of different numbers of proximal and tuft dendrites and the corresponding EPSPs under control conditions where only leak current is present **(E)**, leak + Ka **(F)**, leak + kir **(G)**, and leak x2 **(H)**. Summary values depicted as mean ± SEM

### Rescue of synaptic integration and plasticity

The aLTP, theta burst LTP, and dendritic integration results from Figures 2 and 3 suggest that, given enough synaptic input, *Dlg2+/−* hets can express LTP despite their reduced input resistance. It follows that by increasing input resistance in the *Dlg2+/−* hets, dendritic integration and aLTP could be effectively rescued. Three separate methods to increase input resistance were tested for their effectiveness in rescuing dendritic integration. The first was the relatively broad-spectrum voltage-sensitive potassium channel blocker, 4-aminopyridine (4-AP)(65), the second was the selective Kv1.3, Kv1.4 blocker CP339818(66) and the third was activation of muscarinic M1 receptors(51, 67). 4-AP caused an increase in input resistance, a reduction in the supra-linearity change point, a trend toward increased maximum duration of the compound EPSP as a ratio to the corresponding slope of the single EPSP, and a repolarisation in resting membrane potential (Fig 8A-E). The effects of 4-AP were not genotype-specific, as there were no drug x genotype interactions. These results support the computational modelling predictions that voltage-sensitive potassium channels attenuate dendritic integration and blocking them facilitates it. Since DLG2 interacts with Kv1.4, the selective blocker CP339818 was used to test whether the upregulation of these channels was responsible for reduced dendritic integration. However, CP339818 had no effect on input resistance or dendritic integration (Supplement 1 Fig S18) indicating that upregulation of these specific A-type potassium channels does not underpin the reduction in dendritic integration in the *Dlg2+/−* hets.

**Figure 8.**
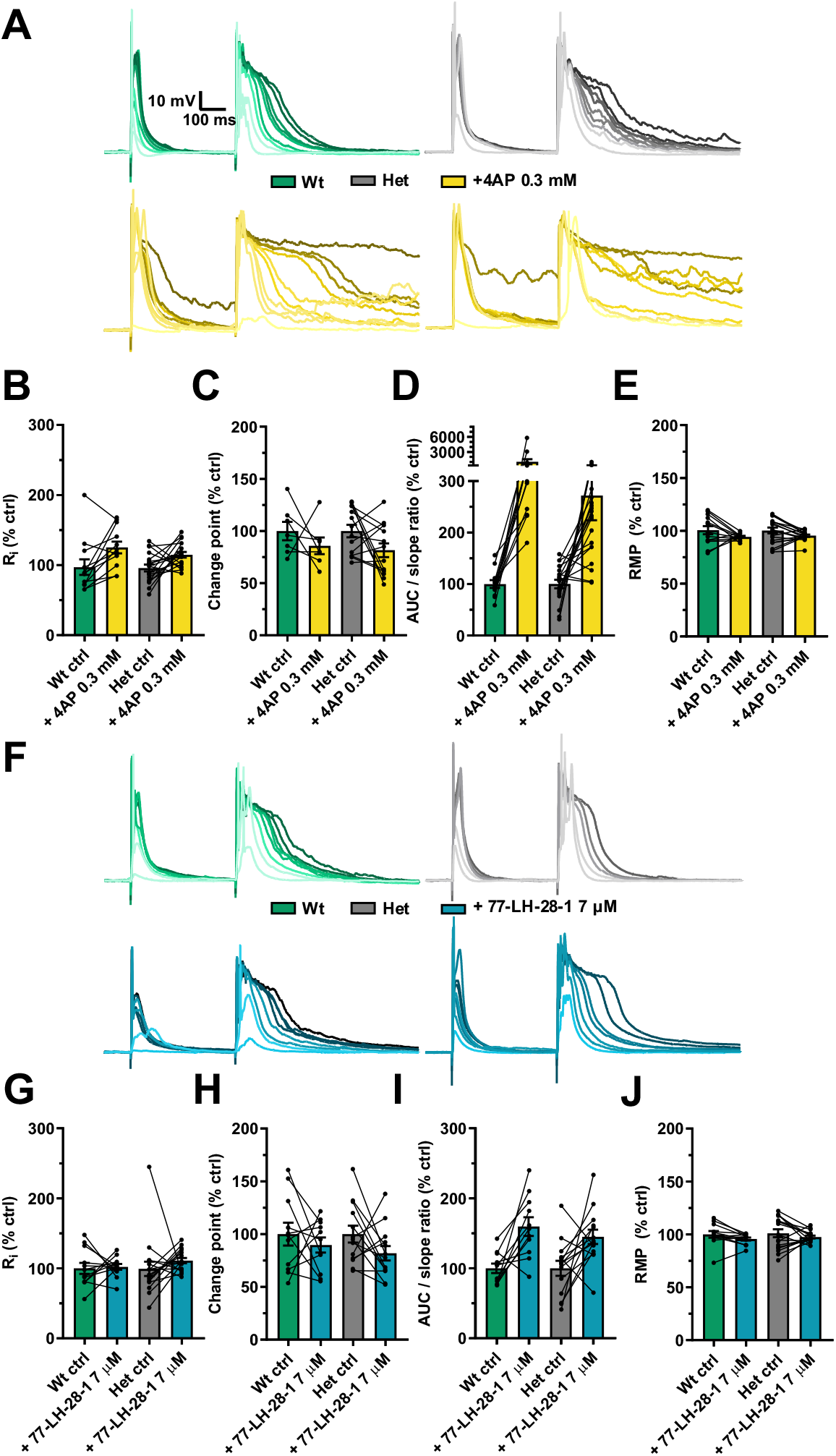
Potassium channel block and M1 agonism lowers dendritic integration thresholds in the *Dlg2+/−* hets. **A)** Example traces depicting a single EPSP followed by a compound EPSP at increasing stimulation intensities (light to dark) over consecutive recording sweeps before and after 4-aminopyridine 0.3 mM across genotype. Input resistance (3-way repeated-measures ANOVA: drug effect: F _1, 23_ = 8.608, P = 0.007, genotype x drug interaction: F _1, 23_ = 0.418, P = 0.524) **(B)**, change point (3-way repeated-measures ANOVA: drug effect: F _1, 14_ = 8.422, P = 0.012, genotype x drug interaction: F _1, 14_ = 0.042, P = 0.841) **(C)**, AUC/slope (3-way repeated-measures ANOVA: drug effect: F _1, 23_ = 3.988, P = 0.058, genotype x drug interaction: F _1, 23_ = 1.570, P = 0.223) **(D)**, and resting membrane potential (RMP) (3-way repeated-measures ANOVA: drug effect: F _1, 24_ = 57.899, P < 0.001, genotype x drug interaction: F _1, 24_ = 0.230, P = 0.636) **(E)** as percent of control before and after the after 4-aminopyridine 0.3 mM across genotype. 32 cells from 15 animals. **F)** Example traces depicting a single EPSP followed by a compound EPSP at increasing stimulation intensities (light to dark) over consecutive recording sweeps before and after 77-LH-28-1 7 µM across genotype. Input resistance (3-way repeated-measures ANOVA: drug effect: F _1, 21_ = 4.12, P = 0.055, genotype x drug interaction: F _1, 21_ = 1.270, P = 0.273) **(G)**, change point (3-way repeated-measures ANOVA: drug effect: F _1, 16_ = 6.879, P = 0.018, genotype x drug interaction: F _1, 16_ = 0.645, P = 0.434) **(H)**, AUC/slope (3-way repeated-measures ANOVA: drug effect: F _1, 17_ = 38.074, P < 0.001, genotype x drug interaction: F _1, 17_ = 0.152, P = 0.701) **(I)**, and resting membrane potential (RMP) (3-way repeated-measures ANOVA: drug effect: F _1, 21_ = 8.931, P = 0.007., genotype x drug interaction: F _1, 21_ = 1.412, P = 0.248) **(J)** as percent of control before and after the after 77-LH-28-1 7 µM. 29 cells from 15 animals. Summary values depicted as mean ± SEM. * P < 0.05, ** P < 0.01, *** P < 0.001 (3-way ANOVA between subject effect)

These results demonstrate, as predicted, that blocking a subset of potassium channels activated around the resting membrane potential facilitates dendritic integration. However, due to the considerable heterogeneity of potassium channels and their ability to compensate for one another coupled with limited availability of selective pharmacological tools, identifying and targeting the precise channels that cause reduced input resistance in the *Dlg2+/−* hets is challenging. An alternative approach, and one with greater therapeutic potential, is to rescue the input resistance reduction indirectly, for example by activation of cholinergic muscarinic M1 receptors that inhibit potassium channel function and increase dendritic excitability(44, 45, 54–58). Support for this approach was found using the highly selective muscarinic M1 receptor allosteric partial agonist 77-LH-28-1(68) which increased input resistance, reduced the change point, increased the maximum duration of the compound EPSP as a ratio to the corresponding slope of the single EPSP, and depolarised the resting membrane potential (Fig 8F-J). However, there were no significant drug x genotype interactions. Similar results were found for the broad-spectrum non-hydrolysable acetylcholine analogue carbachol (Supplement 1 Fig S19).

These results suggest that pharmacological enhancement of dendritic excitability and integration may be sufficient to rescue aLTP in the *Dlg2+/−* hets. Therefore, the aLTP experiment was repeated in the presence of 77-LH-28-1. This rescued aLTP in the *Dlg2+/−* hets with robust aLTP in SC and TA pathways (Fig 9A-C). In addition, unlike in the absence of 77-LH-28-1, there was no effect of genotype and no pathway x genotype interaction (Fig 9C), indicating 77-LH-28-1 selectively rescues aLTP in the *Dlg2+/−* hets. Importantly, baseline EPSC amplitude did not differ among pathways and across genotype (Fig 9E), indicating that the amount of synaptic input received was similar in all conditions. Analysis of the aLTP induction phase revealed that 77-LH-28-1 rescued synaptic summation and the resulting action potential spiking (Fig 9F-H) as well as plateau potential generation (Supplement 1 Fig S20), with the genotypic differences for number of bursts, EPSP summation, and spike number disappearing. Taken together, Figures 8&9 show 77-LH-28-1 reduced the threshold for dendritic integration in both wts and *Dlg2+/−* hets but selectively facilitated aLTP in the *Dlg2+/−* hets indicating induction of synaptic plasticity in the *Dlg2+/−* hets is more sensitive to increased dendritic excitability and synaptic integration.

**Figure 9.**
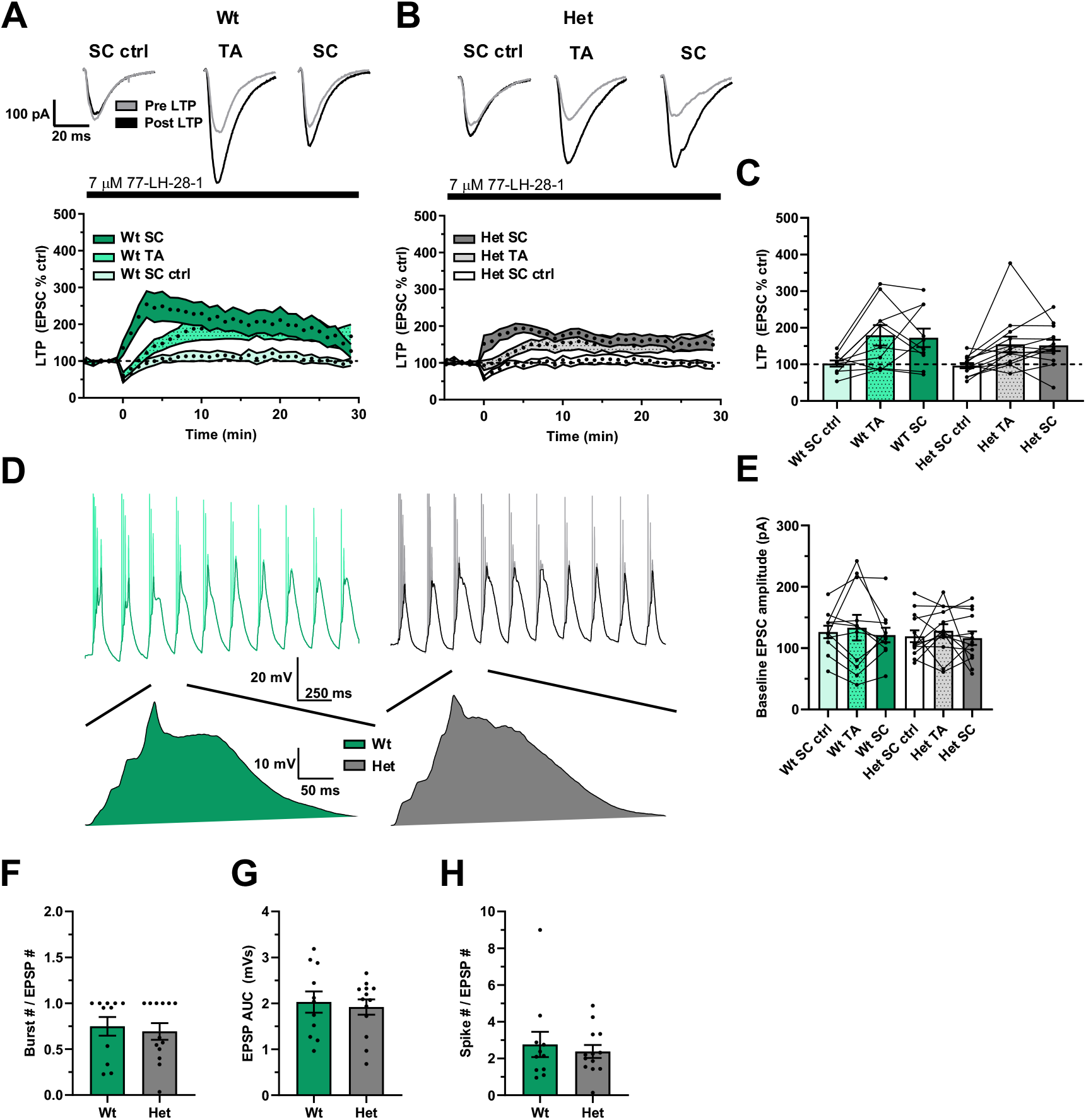
Muscarinic M1 agonism rescues aLTP in the *Dlg2+/−* hets. aLTP over time in wts **(A)** and *Dlg2+/−* hets **(B)**. Example traces pre- and post-induction are displayed for the wt and het groups above their corresponding plots of LTP over time. **C)** aLTP at the 25-30 minute mark post induction across genotype (3-way repeated-measures ANOVA: pathway effect: F _2, 32_ = 12.169, P < 0.001. Genotype main effect: F _1, 16_ = 0.176, P = 0.680. Genotype x pathway interaction: F _2, 32_ = 0.09, P = 0.914). **D)** Example traces of LTP induction, with example EPSPs following *post hoc* spike truncation **E)** Baseline EPSC amplitude across genotype (3-way repeated-measures ANOVA: pathway effect: F _2, 32_ = 1.297, P = 0.287. Genotype main effect: F _1, 16_ < 0.001, P = 0.985. Genotype x pathway interaction: F _2, 32_ = 0.043, P = 0.958). Burst number (3-way ANOVA: genotype main effect: F _1, 16_ = 0.001, P = 0.974) **(F)**, EPSP AUC (3-way ANOVA: genotype main effect: F _1, 16_ = 0.100, P = 0.755) **(G**), and total spike number (3-way ANOVA: genotype main effect: F _1, 16_ = 0.029, P = 0.867) **(H)** across genotype during LTP induction. Summary values depicted as mean ± SEM. * P < 0.05, ** P < 0.01, *** P < 0.001 (3-way ANOVA between subject effect)

## Discussion

NMDAR currents are increased in the *Dlg2+/−* heterozygous rat model. Additionally, dendritic arborisation is reduced. These observations would be expected to combine to enhance neuronal excitability, dendritic integration and synaptic plasticity. Instead, the effects are entirely offset, and indeed reversed, by a concomitant reduction in input resistance caused by an increase in potassium channel expression, most likely A-type potassium channels. This increase in electrical leak is the dominant effect, resulting in a final phenotype where dendritic integration and aLTP are impaired. Crucially, dendritic integration can be rescued by potassium channel block or activation of muscarinic M1 receptors, the latter of which can also rescue synaptic plasticity. These phenotypes are potentially particularly relevant since the *Dlg2+/−* rat model relates to human single copy genetic variants.

The direct interaction between DLG2 and GluN2b NMDAR subunits suggest the most important effects of DLG2 perturbations are on NMDAR function – synaptic integration and plasticity. However, previous studies on *Dlg2−/−* full knockout models have either reported no changes in the AMPA/NMDA ratio or a reduction in the AMPA/NMDA ratio due to reduction in AMPAR function(32–34). Here, AMPAR function was unchanged and instead we found an unexpected increase in NMDAR currents, likely caused by increased expression. On its own, this finding predicts enhanced aLTP, but we found the converse with aLTP impairment. This is similar to previous reports in *Dlg2−/−* mice but with important differences. In homozygous *Dlg2−/−* mice CA1 LTP was reduced in response to the strong TBS induction protocol(33), but in our study using heterozygous *Dlg2+/−* rats TBS-induced LTP was normal. An LTP deficit only became apparent in the *Dlg2+/−* model when neurons were required to integrate converging inputs suggesting a more nuanced but potentially more behaviourally relevant phenotype in the clinically relevant *Dlg2+/−* model. Furthermore, synaptic integration and the initiation of non-linear dendritic events are key determinants of feature detection and selectivity in neuronal networks(46, 69, 70) and a deficit in detecting events and giving appropriate salience are important features of many psychiatric disorders(71).

The dichotomy between enhanced NMDA currents and reduced NMDAR function in *Dlg2+/−* rats during aLTP highlights the dominant role played by changes to intrinsic neuronal excitability; in this instance reduced input resistance caused by increases in potassium channel function. Interestingly, in a *Dlg2−/−* full knockout model no changes in input resistance were reported(72) highlighting again the importance of using clinically relevant models. In our *Dlg2+/−* model this increase in potassium channel function does not appear to be caused by a direct interaction with DLG2 but instead as a homeostatic regulatory mechanism perhaps to compensate for increased synaptic currents. A similar compensatory mechanism is found in other models of psychiatric disorders such as *Fmr1-/y* mice where changes in intrinsic neuronal excitability dominate the resulting perturbations in network processing including dendritic integration and synaptic plasticity(60, 73, 74). This raises the intriguing possibility that genetic disruptions to synaptic function may generally cause homeostatic compensations in intrinsic neuronal excitability that dominate neuronal function and present a common biological phenotype across multiple psychiatric disorders(75).

We have demonstrated in this study that the compensatory mechanisms affecting neuronal excitability can be ameliorated pharmacologically with the administration of selective agonists such as 77-LH-28-1 rescuing impairments in synaptic integration and plasticity, a proof of principle that may be applicable to other psychiatric disorder risk variants. For example, an increase in input resistance due to the administration of 77-LH-28-1 could facilitate spike backpropagation, potentially rescuing the plasticity impairment reported in the *Cacna1c+/−* model of genetic vulnerability to schizophrenia(76). Highly selective muscarinic M1 receptor agonists have efficacy clinically with negligible side effects(77–80) making them attractive pharmaceutical tools. It remains to be seen whether behavioural impairments in DLG2 models can be rescued using similar pharmacological strategies.

## Acknowledgements

We thank Jenny Carter for coordinating the initial generation and breeding of the Dlg2+/− rat line, Rachel Humphries for computational modelling discussions and Aleks Domanski and all members of the Robinson and Mellor groups for general discussions. We also thank Hannah Jones and Estela Michail for their input in the study of morphology. The authors gratefully acknowledge funding from Medical Research Council (UK) (CO’D, PhD studentship to SG), Biotechnology and Biological Science Research Council (UK) (JRM), Wellcome Trust (UK) (JRM, PhD studentship to SW). The Dlg2+/− rats were generated as part of a Wellcome Trust Strategic Award ‘DEFINE’ (JH and LSW) and the Wellcome Trust Strategic Award and the Neuroscience and Mental Health Research Institute, Cardiff University, UK provided core support.

The data in this study was presented as poster and abstract at FENS (2020) and BAP (2021).

## Disclosures

The authors declare no conflict of interest. ER has received research funding from Boehringer Ingelheim, Eli Lilly, Pfizer, Small Pharma Ltd. and MSD, and DD has received research funding from Eli Lilly, but these companies were not associated with the data presented in this manuscript.

