## Supplement 1 for "Reduced expression of the psychiatric risk gene DLG2 (PSD93) impairs hippocampal synaptic integration and plasticity"

**Contents**

*CRISPR-Cas9 generation of Dlg2+/- heterozygous rat model and quality control measures (including Figures S1, S2 and Table S1).*

*Methods: brain slice preparation, electrophysiology, protein quantification and computational modelling*

*Table S2 Peak channel conductances used in computational modelling simulations*

*Table S3 Model parameters used in computational modelling simulations*

*Figure S3 mEPSC NBQX*

*Figure S4 Pathway check*

*Figure S5 Paired pulse facilitation (AMPA/NMDA ratio experiment)*

*Figure S6 aLTP in dorsal-ventral hippocampus*

*Figure S7 Spikes and AUC correlated with aLTP*

*Figure S8 Increased baseline EPSC amplitude produces aLTP in both genotypes*

*Figure S9 Increased baseline EPSC amplitude facilitates plateau potentials and aLTP*

*Figure S10 Theta burst LTP in dorsal-ventral hippocampus*

*Figure S11 Effect of genotype on impedance and resonant frequency*

*Figure S12 Impedance and resonant frequency in dorsal-ventral hippocampus*

*Figure S13 Impedance and resonant frequency across sex*

*Figure S14 Effect of genotype on additional intrinsic properties (rheobase experiment)*

*Figure S15 Intrinsic properties in dorsal-ventral hippocampus (rheobase experiment)*

*Figure S16 Intrinsic properties across sex (rheobase experiment)*

*Figure S17 Computational modelling simulations over entire dendritic arbor*

*Figure S18 Effects of Kv1.3 and Kv1.4 block on dendritic integration*

*Figure S19 Effects of low-dose carbachol on dendritic integration*

*Figure S20* Muscarinic M1 agonism facilitates plateau potential generation in aLTP induction

*References*

*CRISPR-Cas9 generation of Dlg2 heterozygous rat model and quality control measures*.

The *Dlg2*+/- heterozygous rat model was created by Horizon Discovery (St Louis, Missouri, USA). Proprietary bioinformatics software (Horizon Discovery, St. Louis, USA) was used to design a short guide RNA (sgRNA - CCAGGGTCATCTCCAATGTGagg) targeting a Protospacer Adjacent Motif (PAM) sequence within exon 5 of the rat Dlg2 gene on chromosome 1. As shown in Fig S1 the targeting was predicted to result in a 7bp deletion (782933-782939 in the genomic sequence) and consequent downstream frame shift in exon 6 leading to a premature stop codon.

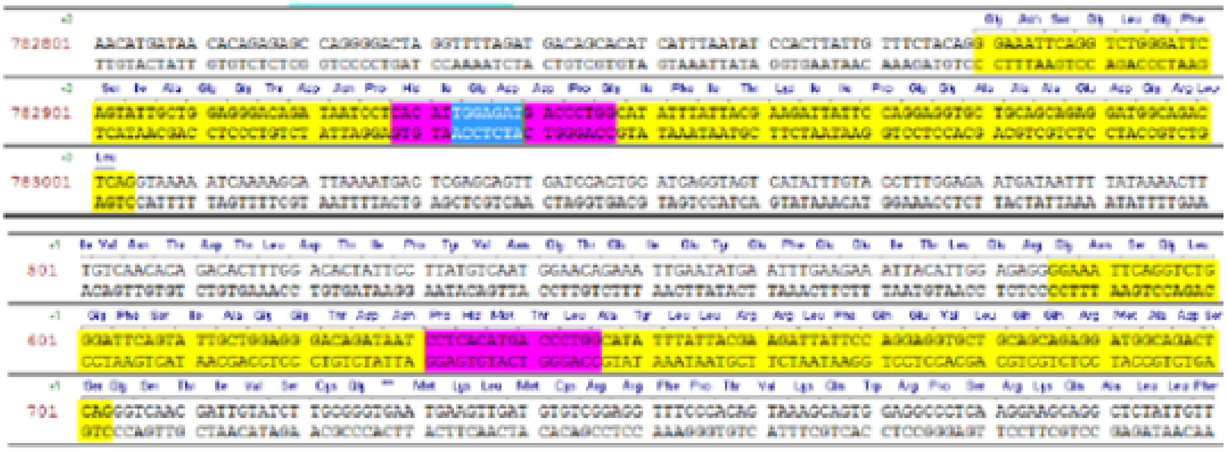

Figure S1: Targeting of exon 5 with deletion shown in blue (top panel) and outcome after non-homologous end joining in purple (lower panel)

Prior to the generation of the founder rats an initial in-vitro confirmation of the efficiency of the sgRNA-Cas9 was demonstrated by nucleofecting the sgRNA-Cas9 into rat C6 glial cells. Genomic DNA (gDNA) PCR products were subsequently generated from nucleofected C6 cells using primers flanking the sgRNA site. gDNA PCR products were screened for non-homologous end joining activity and deletion mutations using the SURVEYOR Cel-1 Mutation Detection Assay (Integrated DNA Technologies). Founder rats were then produced as follows; embryo donor female Long Evans rats were super-ovulated with pregnant mare serum (PMS) and given human chorionic gonadotrophin (HCG) 48 hrs post PMS administration. Females were immediately mated to stud males after HCG administration. Embryo donor females were euthanized 18-24 hrs after mating and their one-cell fertilized embryos were isolated by harvesting the reproductive tract and rupturing the ampulae. Harvested embryos were put in culture media in a CO2 incubator until ready for microinjection. One-cell stage embryos were microinjected with the validated sgRNA-Cas9 and then implanted into synchronized pseudo-pregnant Long Evans recipient females.

The success of the targeting strategy was confirmed in founder rats using sequencing of gDNA PCR products derived from P14 tissue biopsies. Fig S2 illustrates genomic sequencing of a Dlg2 7bp out of frame heterozygous deletion founder. Manual reading of each double peak in the sequencing chromatograph (ABI Sequence Scanner) shows that upstream of the deletion, sequences from the wild-type and modified allele are identical. At the site of the deletion i) the sequence read becomes mixed and ii) the sequence of the secondary peaks from the modified allele align with wild-type sequence, except they occur 7bp further upstream revealing the size and position of the modified allele. As detailed in Fig 1, the effects of the predicted downstream premature stop codon in exon 6 was confirmed by Dlg2 expression analysis which showed the anticipated c.50% reduction in Dlg2 expression in the heterozygotes.

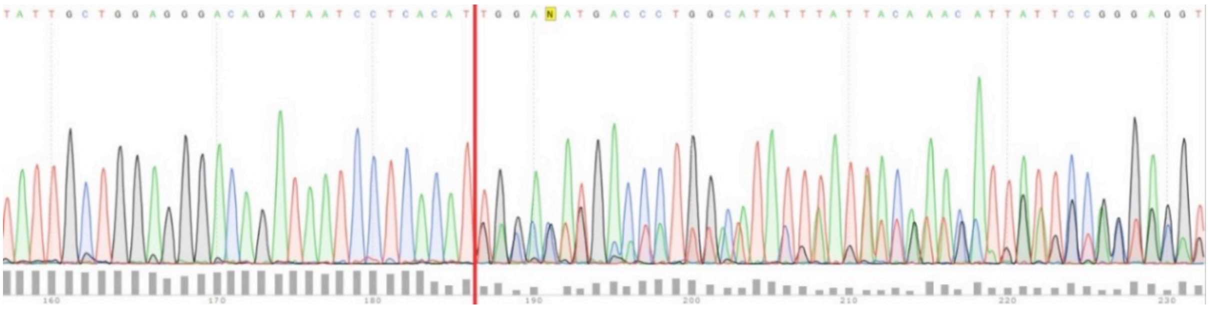

Figure S2: Sequencing of gDNA PCR products from wild-type and modified allele. The deletion site is marked by a red line, prior to which single peaks are seen and after which the peaks in the modified allele show the effects of the deletion moving them 7bp upstream. Data provided by Horizon Discovery and analysed using Snapgene Viewer 3.3.2.

Possible off-target hits were assessed by generating a list of top 10 potential off-target (OT) sites, based on the sgRNA sequence used (CCAGGGTCATCTCCAATGTGagg) and ranked using the MIT website http://crispr.mit.edu/. The top 10 OT sites were computed by taking into account the following i) total number of mismatches, ii) mismatch absolute position (to accommodate for the relatively high disturbance of mismatches falling close to the PAM site) and iii) mean pairwise distance between mismatches to account for the steric effect of closely neighbouring mismatches in disrupting sgRNA-DNA interaction. Corresponding PCR primer pairs were designed to flank the top 10 potential off target sites. Using extracted gDNA from the founder animal and wild-type controls, gDNA products were generated to flank each potential OT site (~300-500bp amplicon) and run on the SURVEYOR assay. Confirming the specificity of the CRISPR-Cas9 targeting to exon 5 of the Dlg2 gene none of the 10 OT sites tested revealed NHEJ activity (Table S1).

Injected sgRNA sequence: CCAGGGTCATCTCCAATGTGagg

OT Site 1 CCAGGGTCATCTCCAATGTTAAG chr19: 34563666-34563688 screened by SURVEYOR aggttagtccgtgcatggtg gcttgcctacagccgtattt - negative

OT Site 2 CTTGAGTCATCTCCAATGTGTGG chr11: 75326973-75326995 screened by SURVEYOR tgtcagtgctgccttttgtc tcctttgtgtggtgtggttc - negative

OT Site 3 CCAGGGTGATCTCCAATCTGAGG chr14: 84150501-84150523 screened by SURVEYOR ggtaactggcctttgggttt tctgatttggggcttaggtg - negative

OT Site 4 TTATTGTCATCTCCAATGTGTAG chr16: 22863405-22863427 screened by SURVEYOR tacccacttttcacccaagc ttgccccttcagagaagac - negative

OT Site 5 CTATGGTCATCTCCAATGGGTAG chr20: 16439248-16439270 screened by SURVEYOR aaaccggttatgtcctgtgc ggaggaagatggagggaaac - negative

OT Site 6 ACAAGCTTATCTCCAATGTGTAG chr6: 46008366-46008388 screened by SURVEYOR tgcctaggaaactggcaact tgtgtcacttggatggatgtc - negative

OT Site 7 CCAGGATAATATCCAATGTGTAG chr16: 34645325-34645347 screened by SURVEYOR tgctcactgctgataggtctg ggatatcattggaccccaca - negative

OT Site 8 TTAGGGTAATATCCAATGTGAGG chr3: 89806272-89806294 screened by SURVEYOR tatctcggcccaagaagaag caaagaccaggatcccaatg - negative

OT Site 9 CAATAATCATCTCCAATGTGCAG chr4: 164282994-164283016 screened by SURVEYOR agcaggtcttcagcttggtt ccagaggccctcaaattaca - negative

OT Site10 CGAAGGTCCACTCCAATGTGCAG chr17: 19811676-19811698 screened by SURVEYOR tcgtgggaaggaaagacttg ggcagtccatgcctgtttat – negative

Table S1: List of top 10 most likely off-target sites (OT) when employing the sgRNA used to make the 7bp deletion in exon 5 of the Dlg2 gene generated using the in-silico MIT online tool (http://crispr.mit.edu/). As shown, gDNA PCR products amplified from the 10 possible off-target sites were all negative (i.e. showed no evidence of deletion or any other genomic change) when screened by the SURVEYOR assay. Data provided by Horizon Discovery.

Founder rats were mated with wild types at Horizon Discovery and generated F2 progeny containing the mutation, thus confirming germ-line transmission. A total of five male positives were exported to Charles River, Lyon, France for re-derivation by embryo transfer. The resulting specific pathogen free (SPF) progeny were sent to Charles River, Margate, UK for routine breeding and maintenance of the lines. The standard breeding protocol was a heterozygous x wild-type cross giving rise to 1:1 average *Dlg2+/-* /wild-type progeny allowing full use of the litter and the generation of littermate controls. The *Dlg2+/-* rat model is viable and Charles River has reported no adverse effects on breeding performance, development, general health and in addition, no deviation from the expected Medelian 1:1 ratio of Dlg2+/- to wild-types and no skewing of the sex ratios. For the current experiments a cohort of breeders was transported to University of Bristol Animal Service Unit and the experimental animals used in the work were generated using heterozygous x wild-type cross giving rise to 1:1 average *Dlg2+/-* /wild-type littermate control progeny.

*Methods: brain slice preparation, electrophysiology, protein quantification and computational modelling*

*Brain slice preparation*

Rats were killed, the brains removed, and hippocampi dissected and sliced in ice-cold sucrose-based solution containing (in mM): 205 Sucrose, 10 Glucose, 26 $NaHCO_3$, 2.5 KCl, 1.25 $NaH_2PO_4$, 0.5 $CaCl_2$, 5 $MgSO_4$, saturated with 95% $O_2$ and 5% $CO_2$. Transverse 400 µm hippocampal slices were cut using the Leica LS1200 vibratome. Slices were incubated in artificial cerebrospinal fluid (aCSF) at 35 °C for 30 min and then at room temperature for 30 min. aCSF contained (in mM): mM: 124 NaCl, 3 KCl, 24 $NaHCO_3$, 1.25 $NaH_2PO_4$ 10 Glucose, 2.5 $CaCl_2$, 1.3 $MgSO_4$, saturated with 95% $O_2$ and 5% $CO_2$.

*Whole-cell patch-clamp recordings*

The recording chamber was perfused with oxygenated aCSF at 32 °C. Slices were visualised using a differential interference contrast Scientifica SliceScope microscope. Borosilicate glass pipettes (pipette resistance of 4-7 MΩ) were pulled using a horizontal P-97 Sutter-instruments puller. 3 different internal solutions were used, potassium-based, potassium-based with Qx-314, and caesium-based. They contained (in mM):

Potassium-based: 120 KMeSO3, 8 NaCl, 10 HEPES, 4 Mg-ATP, 0.3 Na-GTP, 0.2 EGTA, 10 KCl, ~295 mOsm, 7.4 pH

Potassium-based with QX-314: 120 KMeSO3, 8 NaCl, 10 HEPES, 4 Mg-ATP, 0.3 Na-GTP, 0.2 EGTA, 10 KCl, 1 QX-314Cl, ~295 mOsm, 7.4 pH

Caesium-based: 130 CsMeSO3, 4 NaCl, 10 HEPES, 0.5 EGTA, 10 TEA, 2 Mg-ATP, 0.5 Na-GTP, 1 QX-314Cl, ~290 mOsm, 7.4 pH.

Neurobiotin (1 mg/mL) was added to the internal solution used in some intrinsic properties experiments for post hoc cell morphology investigation.

Recordings were obtained using a Multiclamp 700A Molecular Devices amplifier, filtered at 2.4 kHz, and sampled at 10, 20, or 25 kHz, depending on experiment, using a Cambridge Electronic Designs Micro 1401 data acquisition board. Cambridge Electronic Designs Signal

5.12 and Spike 2.7 software were used for data acquisition. All data are presented without adjustment for the junction potential (~−15 mV).

*Synaptic stimulation and plasticity protocols*

All recordings were made in the presence of picrotoxin 50 μM. Schaffer collateral (SC) and temporoammonic (TA) fibres were alternatively stimulated using a paired pulse protocol.

*Experiment: AMPA:NMDA ratio*

Using the caesium-based internal solution, cells were held at -70 mV for 10 min, +40 mV for 5 min, and -70 mV for another 10 min. This allowed for the acquisition of predominantly AMPA-mediated EPSCs at -70 mV and combined AMPA- and NMDA-mediated EPSCs at +40 mV. AMPA-mediated EPSCs were measured at the peak amplitude, whilst NMDA-mediated EPSCs were measured at 45 ms after the first stimulation artefact. If the AMPA-mediated EPSCs recorded before and after the voltage switch differed by more than 30%, data was excluded from the analysis.

*Experiment: AMPA mEPSC*

mEPSCs were recorded using the caesium-based internal solution. Recordings were made in the presence of TTX 500 nM and at a holding potential of -65 mV. 10 μM NBQX was applied in a subset of the initial experiments to confirm that the mEPSCs were AMPA-mediated (Fig S2). The following parameters were used for mEPSC detection post hoc: -3 pA amplitude, 0.1 ms tau (rise), 3 ms tau (decay), dead time 7 ms, rising edge window 2 ms.

*Experiment: SK-mediated modulation of NMDAR*

With a potassium-based QX-314 internal solution, cells were held in current-clamp with enough current injection to keep them at approximately -50 mV. The aCSF contained CGP55845 1μM throughout the experiment, with subsequent applications of apamin 100 nM alone and together with and D-APV 50 μM for 10 min each. Compound EPSPs were elicited using a burst of 5 stimulations at 100 Hz. EPSP decay was modelled using a single exponential function fit between the linear component of the decay of the final EPSP and baseline, from the averaged traces of the final 3 min of every condition.

*Experiment: GluN2b-mediated EPSC decay*

With a caesium-based internal solution, cells were held at +40 mV. aCSF contained NBQX 10 μM throughout the experiment, with subsequent additions of RO256981 1 μM alone and in combination with D-APV 50 μM for 10 min each. The decay of the second EPSC was modelled using a double exponential function fit between the amplitude peak and the baseline, from the averaged traces of the final 3 min of every condition.

*Experiment: Paired theta burst LTP in SC synapses*

Using a potassium-based internal solution, SC and TA fibres were alternatively stimulated in the voltage clamp configuration at -65 mV. With a 5 min baseline and within 10 min of breaking into the whole-cell configuration, LTP was induced in the SC synapses using a 2 second paired theta burst (5 Hz) protocol with simultaneous presynaptic stimulation and somatic depolarization steps. LTP induction was performed in the current-clamp configuration with enough current injection to maintain the cell at -65 mV. LTP was measured at 25-30 min post induction, using the TA pathway as a negative control.

*Experiment: Associative LTP in* SC *and TA synapses*

Using a potassium-based internal solution, the cell was held at -65 mV. SC (from either side of the recording cell) and TA fibres were alternatively stimulated. With a 5 min baseline and within 10 min of breaking into the whole-cell configuration, LTP was induced in the one of the SC and the TA synapses using a 2 s theta burst (5 Hz) protocol with simultaneous presynaptic stimulation of the SC and the TA fibres, with no somatic depolarization steps. The SC pathway from the other side of the recording cell (the one that did not participate in LTP induction) was used as a control pathway. The cell was maintained at -55 mV in current clamp during induction and the theta burst train was repeated for a total of 3 times, at a 20 s interval. Initial baseline EPSC amplitudes were adjusted to be approximately 100 pA. LTP was measured at 25-30 min post induction. A pathway check was run at the end of the experiment to ensure the synapses stimulated belonged to distinct pathways (Fig S3). The experiment was repeated in the presence of 77-LH-28-1 7 μM, with baseline EPSC amplitudes adjusted to be approximately 100 pA.

*Experiment: Supralinear integration of dendritic spiking into plateau potentials*

Using a potassium-based internal solution containing QX-314, a single and a compound EPSP (burst of 5 stimulations at 100 Hz) in the SC pathway were recorded using increasing stimulus intensity in the presence of CGP55845 1µM. The subthreshold rising EPSP slope of the single EPSP was used to normalise area under the curve comparisons of the compound EPSP across conditions. The area under the curve was calculated from 50 ms to 450 ms after the compound EPSP. The slope-area relationship was smoothed using Savitzky–Golay filtering. The data was then passed through a change point analysis function in Matlab designed to find changes in signal to identify the threshold of nonlinearity (1). The maximum recorded area under the curve (AUC) and its corresponding single EPSP slope was measured as a ratio to allow between-subject comparisons. D-APV 50 µM was applied at the end of the experiment during ongoing maximal stimulation. The experiment was repeated in the presence of 4AP 0.3 mM, CP339818 5 µM, carbachol 1 µM, or 77-LH-28-1 7 µM.

*Experiment: Intrinsic cell properties*

Using a potassium-based internal solution, sometimes containing 1 mg/mL neurobiotin, intrinsic cell properties were measured in a specific order. Immediately after breaking in whole-cell, the clamping configuration was changed to I=0 to record the resting membrane potential (RMP). Subsequently, the current clamp configuration was adopted for rheobase experiments and enough current was injected to keep the cell at -65 mV. Depolarising 0.8 s current steps of variable magnitude were applied until a single action potential was fired consistently. A chirp current injection followed, 40 pA peak to peak and 20 s in duration, increasing in frequency from 0.2 to 20 Hz, to allow the measurement of impedance and resonance. Subsequently, hyperpolarising and depolarising steps (-150 to 250 pA and 0.8 s). 20 µM ZD-7288 was then washed on, followed by a repeat of the current step and chirp current injection protocols. Impedance was calculated by first applying the fast Fourier transform algorithm to the input current and output voltage sinusoid chirp data and then by taking their complex ratio. Savitzky–Golay filtering was used to smooth the data.

*Immunohistochemistry and morphological analysis*

Brain slices were placed in 4% PFA for 24-40 hours and then kept in phosphate buffered saline (PBS) (4 °C). Slices were washed in PBS, permeabilised in TritonX100 1:100 in PBS for 1 h, washed and incubated in PBS with 3% bovine serum albumin (BSA) 1 h. Alexafluor-594-streptavidin (1:1000) in PBS with 3% BSA was applied to bind to neurobiotin. Slices were washed in PBS with 3% BSA and incubated with DAPI (1:1000) in PBS with 3% BSA. Slices were then mounted onto glass slides. The slices were imaged using a widefield fluorescence microscope, using Leica LAS X software. The acquired images were processed using Fiji imageJ 1.8.0 software. The publicly available Simple Neurite Tracer plugin was used for semi-automated tracing, visualisation, and analysis of the images (2). The Sholl Analysis extension of the Simple Neurite Tracer plugin was also employed (3).

*Protein Quantification*

Western blot analysis was conducted on hippocampal tissue from wt (n=12) and het (n=12) animals. Each tissue sample was lysed in Syn-PER lysis and extraction buffer (Thermo Fisher, UK) with mini protease inhibitor cocktail (Roche Diagnostics) and phosphatase inhibitor (Cell signalling, UK) according to description from manufacturer.  After using BCA Assay kit to measure the total amount of protein in each sample, electrophoresis and blotting were carried out.

Gels (4–12% NuPAGE Bis-Tris Midi, 45 well) were loaded with 40ug of protein was loaded per well. Samples were added to Laemmli buffer at a 1:1 ratio and this mixture heated at 96 $^{o}$C for 5 minutes to denature protein-protein interactions and facilitate antibody bindings. Samples were arranged so that genotypes were counterbalanced across gels with a WT standard used on each gel for comparison. Gels were run at room temperature in NuPAGE$^{TM}$ Running Buffer (Invitrogen, UK) at 85V for 20 mins and then for a further hour at 115V. Protein was then transferred to 0.45 um pore size nitrocellulose membrane (Invitrogen, UK) at 85 V for 2 hours 15 minutes at an ambient temperature of 4 °C in NuPAGE$^{TM}$ Transfer Buffer (Invitrogen, UK) containing 10% 2-propanol (ThermoFisher Scientific, UK). Membranes containing transferred protein were washed in Tris-Buffered Saline (20mM Tris, 150mM NaCl, pH 7.6) with 0.1% Tween 20 (TBST) before blocking in 5% milk for one hour at room temperature with gentle rocking.

Primary antibodies (rabbit anti-PSD93, 1:1000, Cell Signalling Technology, USA; mouse anti-GluN1, 1:1000, Merck Millipore, UK; rabbit anti-PSD95, 1:2000, Abcam, UK; mouse anti-GAPDH, 1:5000, Abcam, UK) were diluted to appropriate concentrations in 5% milk and incubated with the membrane overnight at 4 °C. Membranes were then subject to 3 x 10 minute TBST washes before incubation with the appropriate fluorescent IRDye 680RD secondary antibodies at 1:15,000 dilution in 5% milk at room temperature. After another series of TBST washes membranes were imaged on Odyssey CLx Imaging System (Li-COR, Germany).

Densiometric analysis of bands was performed using ImageLab 6.0 (https://imagej.nih.gov/ij/). The densities (with background subtracted) of the protein of interest were divided by the loading control densities for each sample to provide normalised values. Densities were then averaged by group.

*Computational modelling*

All modelling work was done using the NEURON simulation environment (4). In-house morphological reconstructions of wt and het neurons were used. A total of 6 representative reconstructions (3 wt, 3 het) were populated with mechanisms corresponding to Ka (5-6), Kir (7-9), Km (10), Kdr (5), and NaV (5,10) channels with parameter values tuned to replicate experimental data to assess input resistance in relation to morphology (Table S2). The channel mechanisms were taken from the Neuron Model database (11). To compare the contributions of the different candidate channels to input resistance, a wt reconstruction was populated with leak current and Ka or Kir channel mechanisms, whose conductance was scaled by 0.5-5. To assess the contribution of input resistance and potassium channels on dendritic integration, the following series of steps were performed. Dendrites were assigned a number which was then shuffled (the same seed was used across conditions). Dendrites were cumulatively recruited using a glutamate mechanism (1 synapse per dendrite), with the ratio of apical proximal and apical tuft dendrites systematically iterated upon. This process was then repeated following changes to conditions, such as the inclusion of a potassium channel mechanism. All dendritic integration simulations were done on the representative wt reconstruction. Model parameters are summarised in Table S3 (12-14).

Table S2 peak channel conductances used in computational modelling simulations

| Peak conductance (S/cm$^2$) | | |
|---|---|---|
| **Channel** | Soma | Dendrites |
| **Km** | 0.017 | 0.017 |
| **NaV** | 0.1 | 0.03 |
| **Kdr** | 0.04 | 0.04 |
| **Ka** | 0.02 | 0.02 |
| **Kir** | 0.0000144 | 0.0000144 |
| **Leak (gpas)** | 0.000025 | 0.000025 |

Km M-type potassium channel

NaV Voltage-gated sodium channel

Kdr Delayed-rectifier potassium channel

Ka A-type potassium channel

Kir Inwardly-rectifying potassium channel

Gpas passive conductance

Table S3 model parameters used in computational modelling simulations

| **Model parameters** | |
|---|---|
| **Axial resistivity (Ω cm)** | 150 |
| **Membrane capacitance (µf/cm2)** | 1.5 |
| **Membrane resistivity (kΩ cm2)** | 40 |
| **Resting membrane potential (mV)** | -65 |
| **Temperature (°C)** | 35 |
| **Equilibrium potential (K) (mV)** | -90 |
| **Equilibrium potential (Na) (mV)** | 55 |
| **NMDA rise time constant (ms)** | 4 |
| **NMDA decay time constant (ms)** | 42 |
| **Equilibrium potential (NMDA) (mV)** | 0 |

### Figures

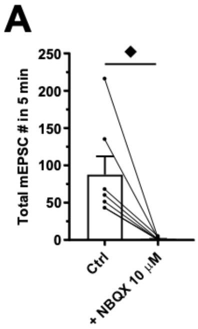


Figure S3 mEPSC are blocked by NBQX. **A)** Total mEPSCs before and after the application of NBQX 10 µM. Summary values depicted as mean ± SEM. ♦ P < 0.05, ♦♦ P < 0.01, ♦♦♦ P < 0.001 (paired t-test)

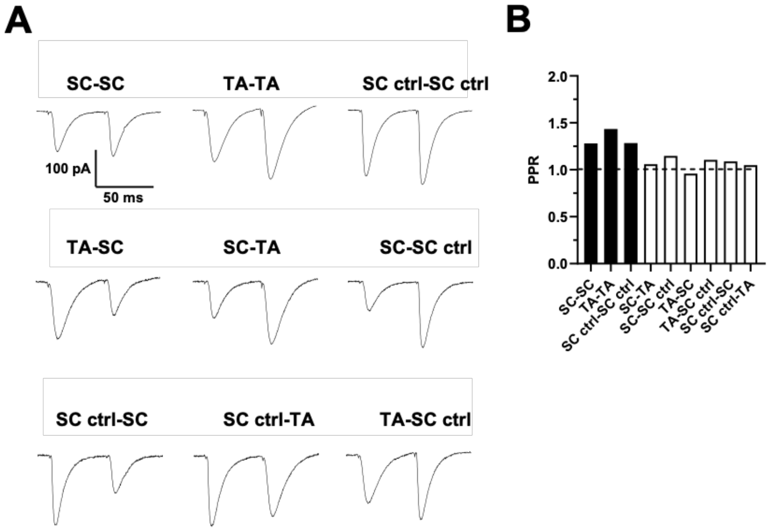


Figure S4 Example pathway independence check from the aLTP experiments. **A)** EPSC traces of 3 pathway responses recorded in different combinations, used to calculate paired-pulse ratio. **B)** Paired-pulse ratios corresponding to the traces in panel A. When the same pathway is stimulated in sequence, there is facilitation. When different pathways are stimulated in sequence, there is little to no facilitation

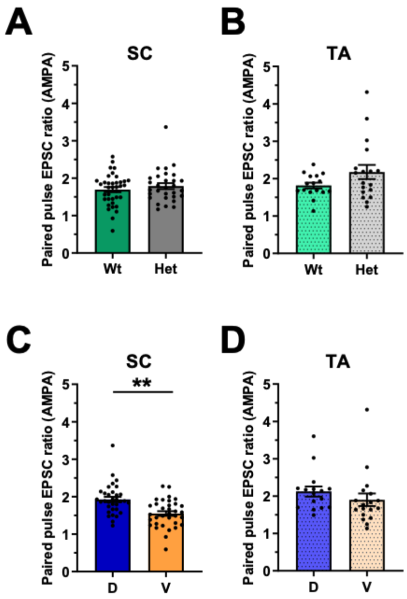


Figure S5 Paired pulse facilitation (AMPA/NMDA ratio experiment). Paired pulse EPSC ratio, taken from the AMPA component at -70 mV, across genotype in the SC (3-way ANOVA: genotype main effect: $F_{1, 67} = 0.010$, $P = 0.921$) **(A)** and TA (3-way ANOVA: genotype main effect: $F_{1, 35} = 1.739$, $P = 0.198$) **(B)** pathways. The same measurement but across dorsal/ventral aspects of the hippocampus in the SC (3-way ANOVA: aspect main effect: $F_{1, 67} = 11.367$, $P = 0.001$) **(C)** and TA (3-way ANOVA: aspect main effect: $F_{1, 35} = 0.087$, $P = 0.770$) **(D)** pathways. 68 cells from 32 animals for the SC data set and 34 cells from 16 animals for the TA data set. Summary values depicted as mean ± SEM. * $P < 0.05$, ** $P < 0.01$, *** $P < 0.001$ (3-way ANOVA between subject effect)

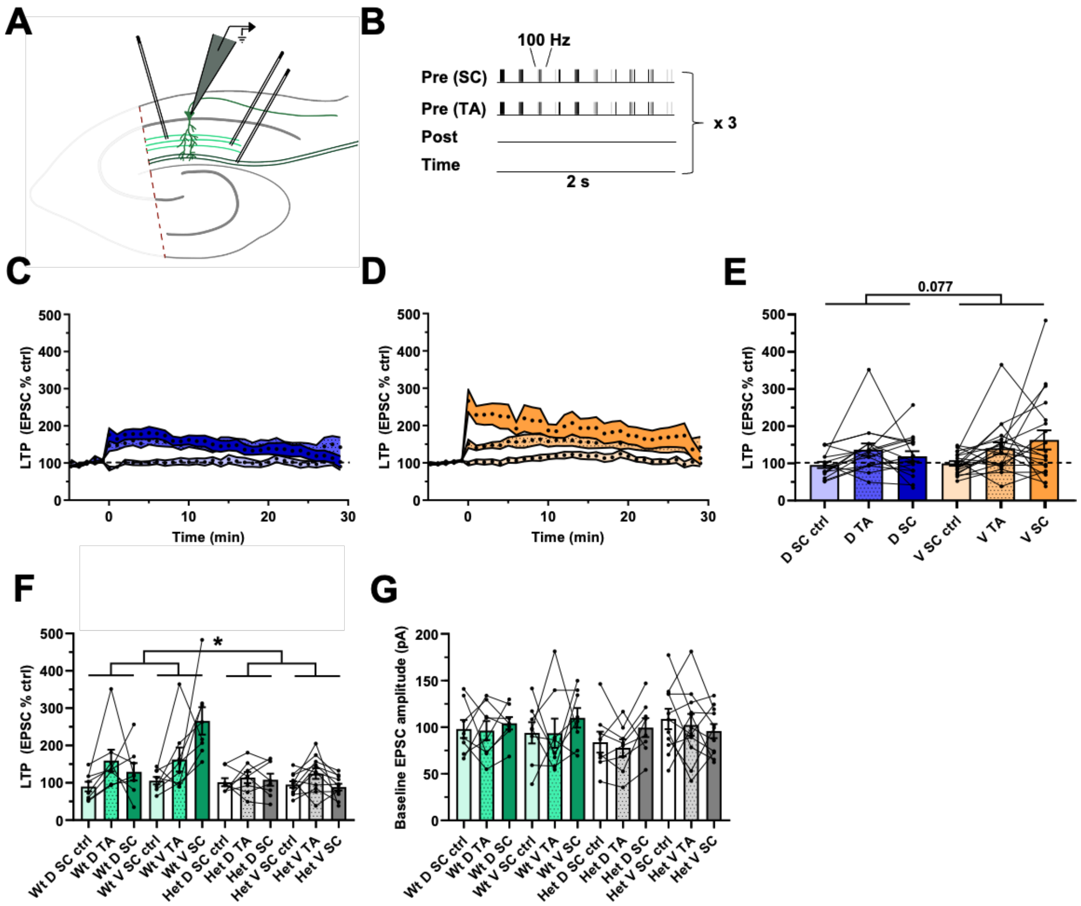


Figure S6 Attenuated aLTP in the dorsal hippocampus. **A)** Schematic representation of the hippocampal slice recording setup, with the CA3 removed and stimulating electrodes in two separate areas of the stratum radiatum and in the stratum lacunosum moleculare. **B)** aLTP induction protocol, where one SC pathway and one TA pathway were tested and where the second SC pathway acted as a negative control. There was no induced somatic depolarisation. This induction protocol was repeated thrice at an interval of 10 seconds. aLTP over time in dorsal **(C)** and ventral **(D)** aspects of the hippocampus. **E)** aLTP at the 25-30 minute mark post induction across dorsal-ventral aspects of the hippocampus and pathways (3-way repeated-measures ANOVA: pathway effect: $F_{2, 54} = 7.300$, $P = 0.002$. Aspect main effect: $F_{1, 27} = 3.374$, $P = 0.077$. Aspect x pathway interaction: $F_{2, 54} = 2.598$, $P = 0.084$). **F)** aLTP at the 25-30 minute mark post induction across genotype, aspects of the hippocampus, and pathway (3-way repeated-measures ANOVA: genotype x aspect interaction: $F_{1, 27} = 4.510$, $P = 0.043$. Genotype x aspect x pathway interaction: $F_{2, 54} = 6.195$, $P = 0.004$). **G)**

Baseline EPSC amplitude across genotype (3-way repeated-measures ANOVA: genotype x aspect interaction: $F_{1, 27} = 1.577$, $P = 0.220$. Genotype x aspect x pathway interaction: $F_{2, 54} = 1.744$, $P = 0.185$). 35 cells from 17 animals. Summary values depicted as mean ± SEM. * $P < 0.05$, ** $P < 0.01$, *** $P < 0.001$ (3-way ANOVA between subject effect)

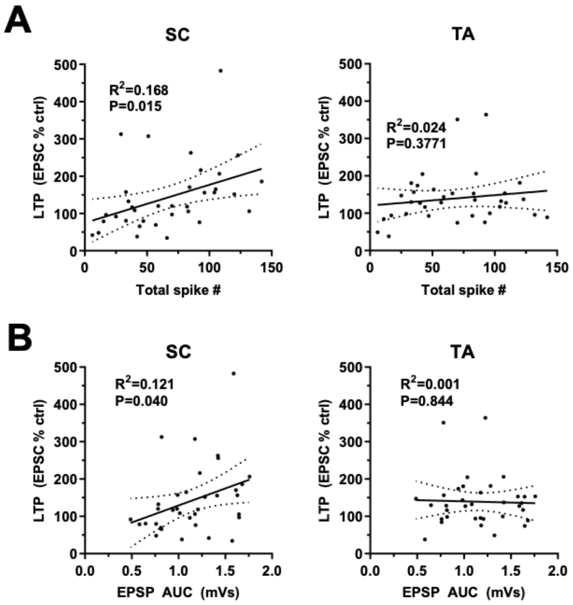


Figure S7 Spikes and AUC are positively correlated with aLTP in the SC but not the TA pathway. Data from the 3-pathway aLTP experiment. **A)** Correlation between total spike number and LTP in the SC (Pearson correlation: $R^2$ = 0.168, P = 0.015) and TA (Pearson correlation: $R^2$ = 0.024, P = 0.377) pathways. **B)** Correlation between EPSP AUC and LTP in the SC (Pearson correlation: $R^2$ = 0.121, P = 0.040) and TA (Pearson correlation: $R^2$ = 0.001, P = 0.844) pathways.

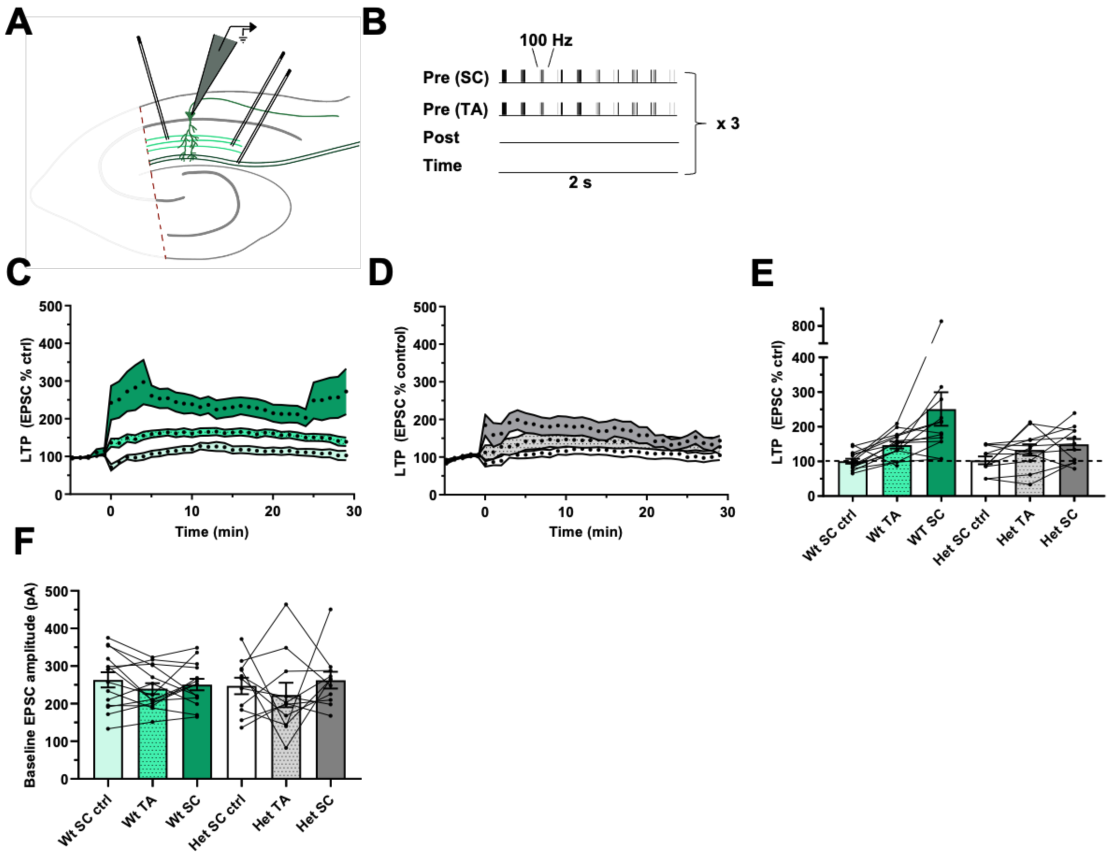


Figure S8 Increased baseline EPSC amplitude produces aLTP in both genotypes. **A)** Schematic representation of the hippocampal slice recording setup, with the CA3 removed and stimulating electrodes in two separate areas of the stratum radiatum and in the stratum lacunosum moleculare. **B)** aLTP induction protocol, where one SC pathway and one TA pathway were tested and where the second SC pathway acted as a negative control. There was no induced somatic depolarisation. This induction protocol was repeated thrice at an interval of 10 seconds. aLTP over time in wts **(C)** and *Dlg2+/-* hets **(D)**. **E)** aLTP at the 25-30 minute mark post induction across genotype and pathway (3-way repeated-measures ANOVA: pathway effect: $F_{2, 36}$ = 6.602, P = 0.004. Genotype main effect: $F_{1, 28}$ = 0.064, P = 0.803. Genotype x pathway interaction: $F_{2, 36}$ = 0.543, P = 0.586). **F)** Baseline EPSC amplitude across genotype (3-way repeated-measures ANOVA: genotype x aspect interaction: $F_{1, 18}$ = 0.144, P = 0.709. Genotype x aspect x pathway interaction: $F_{2, 36}$ = 0.193, P = 0.825). 25 cells from 17 animals. Summary values depicted as mean ± SEM. * P < 0.05, ** P < 0.01, *** P < 0.001 (3-way ANOVA between subject effect)

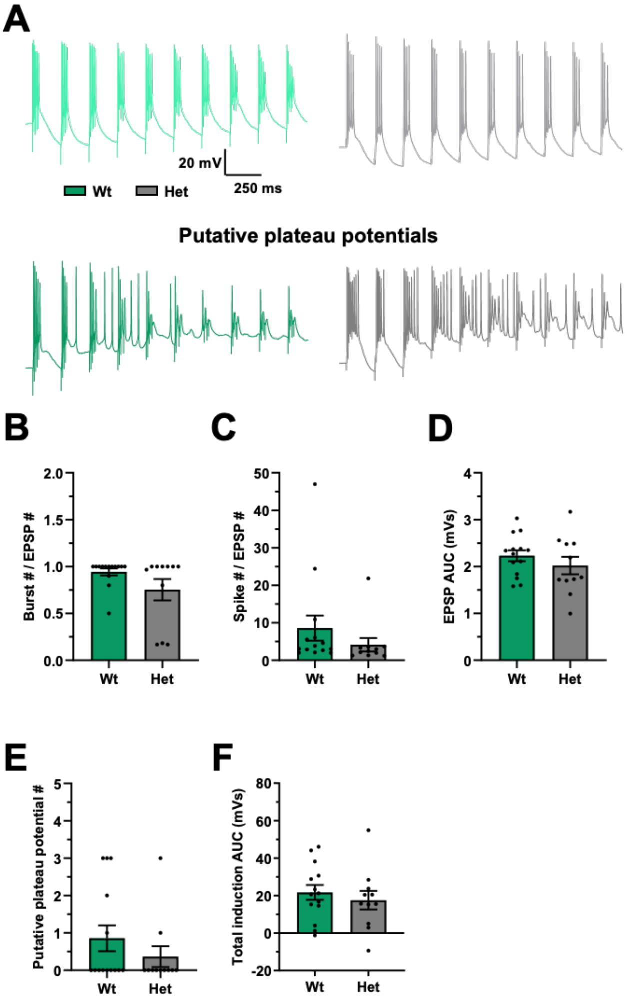


Figure S9 Increased baseline EPSC amplitude facilitates putative plateau potential generation and aLTP. **A)** example aLTP induction traces showing putative plateau potentials. Burst number (3-way ANOVA: genotype main effect: $F_{1, 25} = 1.781$, $P = 0.199$) **(B)**, spike number (3-way ANOVA: genotype main effect: $F_{1, 25} = 0.000$, $P = 0.985$) **(C)**, mean EPSP area under the curve (AUC) (3-way ANOVA: genotype main effect: $F_{1, 25} = 0.478$, $P = 0.498$) **(D)**, putative plateau potential number (3-way ANOVA: genotype main effect: $F_{1, 25} = 0.007$, $P = 0.936$) **(E)**, total induction AUC (3-way ANOVA: genotype main effect: $F_{1, 25} = 0.167$, $P = 0.688$) **(F)** across genotype. 25 cells from 17 animals. Summary values depicted as mean ± SEM. * $P < 0.05$, ** $P < 0.01$, *** $P < 0.001$ (3-way ANOVA between subject effect)

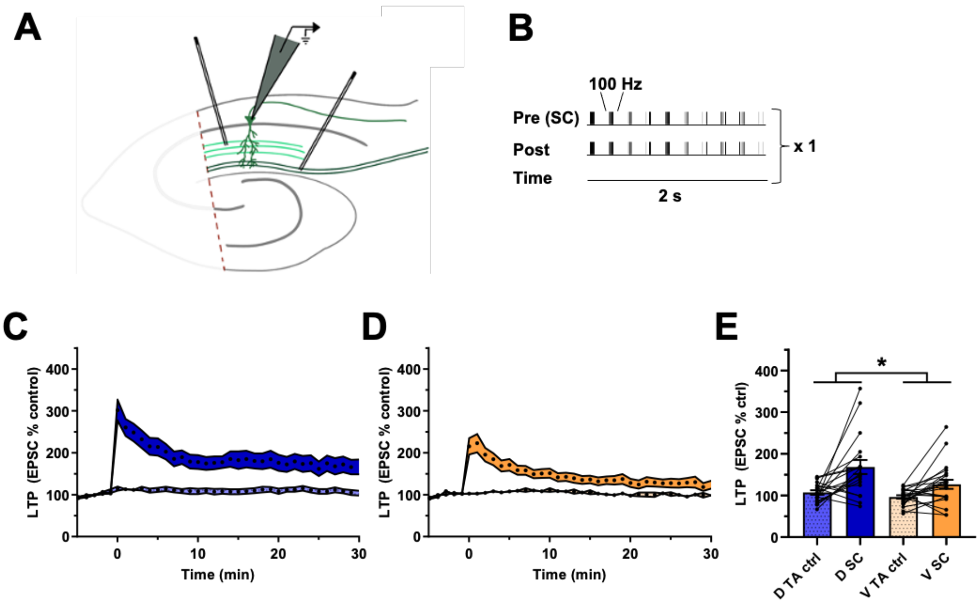


Figure S10 Attenuated theta burst LTP in the ventral aspect of the hippocampus. **A)** Schematic representation of the hippocampal slice recording setup, with the CA3 removed and stimulating electrodes in two separate areas of the stratum radiatum and in the stratum lacunosum moleculare. **B)** Theta burst LTP induction protocol, where the SC pathway was paired with somatic depolarisation and where the TA pathway acted as a negative control. Theta burst LTP over time in doral **(C)** and ventral **(D)** aspects of the hippocampus. **E)** Theta burst LTP at the 25-30 minute mark post induction across genotype (3-way repeated-measures ANOVA: pathway effect: $F_{1, 33} = 18.979$, $P < 0.001$. Aspect main effect: $F_{1, 33} = 5.259$, $P = 0.028$. Aspect x pathway interaction: $F_{1, 33} = 2.085$, $P = 0.158$). 41 cells from 19 animals. Summary values depicted as mean ± SEM. * P < 0.05, ** P < 0.01, *** P < 0.001 (3-way ANOVA between subject effect)

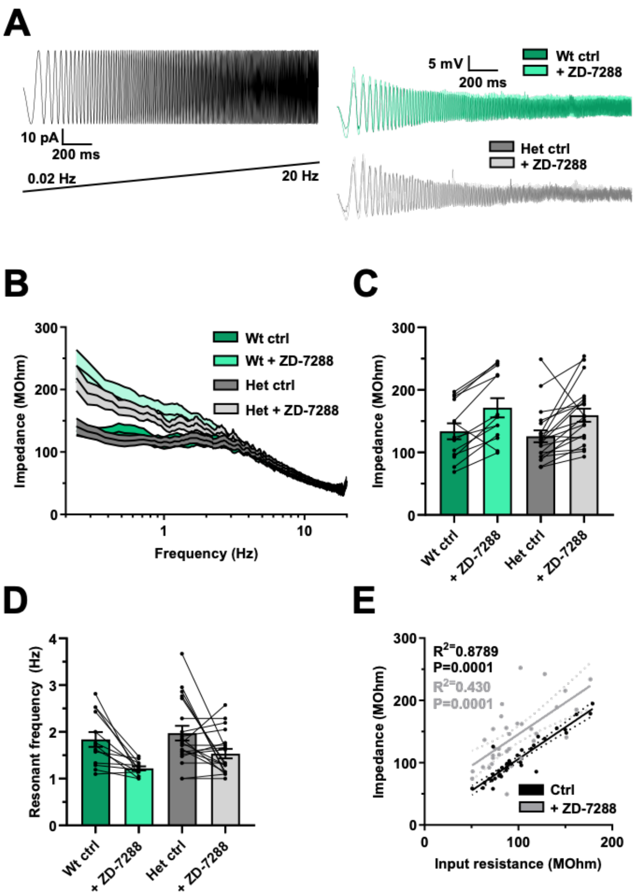


Figure S11 No effect of genotype on impedance and resonant frequency. **A)** input current wave (left) and output voltage waves (right) across genotype and before and after the application of ZD-7288 20 µM. **B)** impedance over 0.2-20 Hz input frequencies before and after the application of ZD-7288 20 µM across genotype. Maximum impedance in the 1-20 Hz range (3-way repeated-measures ANOVA: drug effect: $F_{1, 25} = 26.64$, $P < 0.001$. Genotype main effect: $F_{1, 25} = 0.044$, $P = 0.835$. Genotype x drug interaction: $F_{1, 25} = 0.033$, $P = 0.858$) **(C)** and the corresponding resonant frequency (3-way repeated-measures ANOVA: drug effect: $F_{1, 25} = 22.402$, $P < 0.001$. Genotype main effect: $F_{1, 25} = 1.639$, $P = 0.212$. Genotype x drug interaction: $F_{1, 25} = 1.538$, $P = 0.226$) **(D)**. **E)** Correlation between input resistance and impedance before (Pearson correlation: $R^2 = 0.879$, $P < 0.001$) and after (Pearson correlation: $R^2 = 0.430$, $P < 0.001$) application of ZD-7288 20 µM. 33 cells from 17 animals. Summary values depicted as mean ± SEM. * P < 0.05, ** P < 0.01, *** P < 0.001 (3-way ANOVA between subject effect)

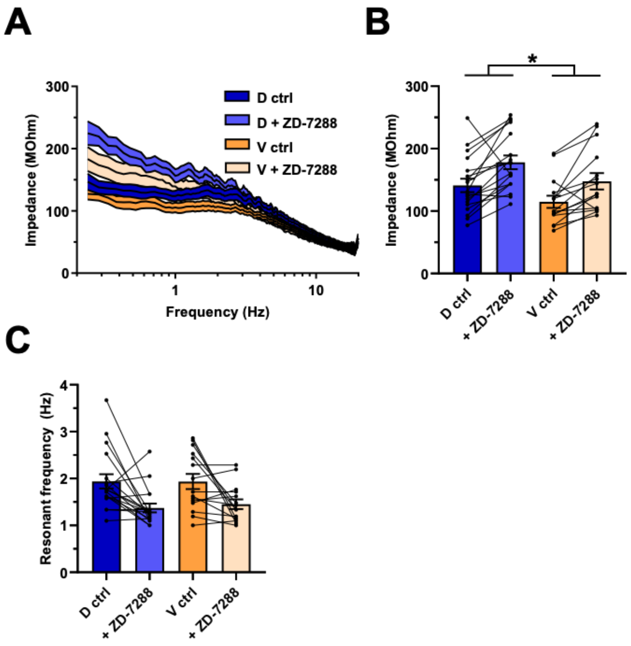


Figure S12 Reduced impedance in the ventral aspect of the hippocampus. **A)** impedance over 0.2-20 Hz input frequencies before and after the application of ZD-7288 20 µM across genotype. Maximum impedance in the 1-20 Hz range (3-way repeated-measures ANOVA: drug effect: $F_{1, 25} = 26.640$, $P < 0.001$. Aspect main effect: $F_{1, 25} = 6.300$, $P = 0.019$. Aspect x drug interaction: $F_{1, 25} = 0.427$, $P = 0.519$) **(B)** and the corresponding resonant frequency (3-way repeated-measures ANOVA: drug effect: $F_{1, 25} = 22.402$, $P < 0.001$. Aspect main effect: $F_{1, 25} = 0.584$, $P = 0.452$. Aspect x drug interaction: $F_{1, 25} = 0.055$, $P = 0.817$) **(C)**. 33 cells from 17 animals. Summary values depicted as mean ± SEM. * $P < 0.05$, ** $P < 0.01$, *** $P < 0.001$ (3-way ANOVA between subject effect)

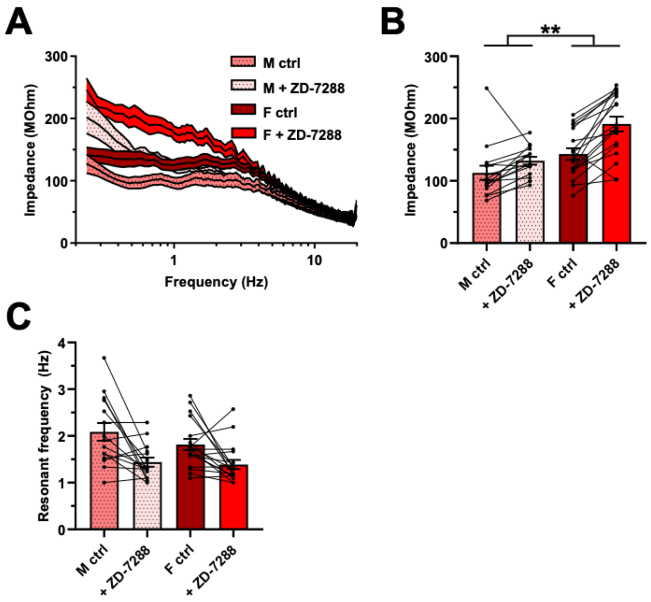


Figure S13 Increased impedance in neurons from female rats. **A)** impedance over 0.2-20 Hz input frequencies before and after the application of ZD-7288 20 µM across genotype. Maximum impedance in the 1-20 Hz range (3-way repeated-measures ANOVA: drug effect: $F_{1, 25} = 26.640$, $P < 0.001$. Sex main effect: $F_{1, 25} = 15.070$, $P = 0.001$. Sex x drug interaction: $F_{1, 25} = 2.943$, $P = 0.099$) **(B)** and the corresponding resonant frequency (3-way repeated-measures ANOVA: drug effect: $F_{1, 25} = 22.402$, $P < 0.001$. Sex main effect: $F_{1, 25} = 1.542$, $P = 0.226$. Sex x drug interaction: $F_{1, 25} = 1.078$, $P < 0.309$) **(C)**. 33 cells from 17 animals. Summary values depicted as mean ± SEM. * $P < 0.05$, ** $P < 0.01$, *** $P < 0.001$ (3-way ANOVA between subject effect)

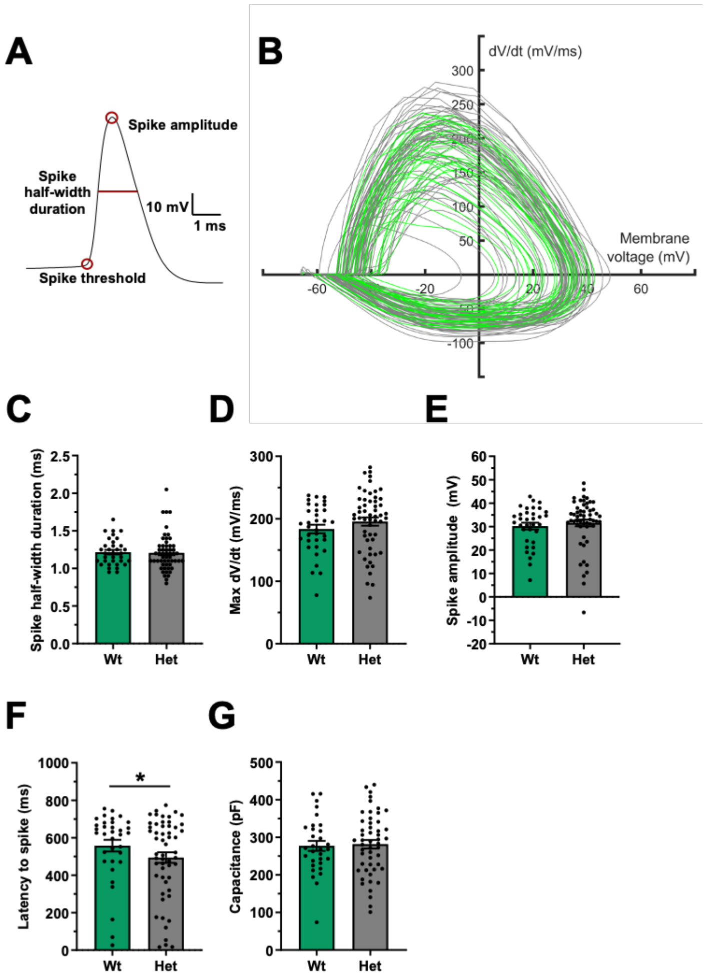


Figure S14 The effect of genotype on additional intrinsic properties. Data from the rheobase set of experiments. **A)** example spike from. **B)** Membrane voltage-spike dV/dT relationship across genotype (wt: green, *Dlg2+/-* het: gray). Spike half-width duration (3-way ANOVA: genotype main effect: $F_{1, 89} = 0.775$, $P = 0.381$) **(C)**, max dV/dt (3-way ANOVA: genotype main effect: $F_{1, 89} = 2.300$, $P = 0.133$) **(D)**, spike amplitude (3-way ANOVA: genotype main effect: $F_{1, 89} = 0.545$, $P = 0.462$) **(E)**, latency to spike (3-way ANOVA: genotype main effect: $F_{1, 89} = 4.122$, $P = 0.046$) **(F)**, and membrane capacitance (3-way ANOVA: genotype main effect: $F_{1, 82} = 0.053$, $P = 0.818$) **(G)** across genotype.136 cells from 38 animals. Summary values depicted as mean ± SEM. * $P < 0.05$, ** $P < 0.01$, *** $P < 0.001$ (3-way ANOVA between subject effect)

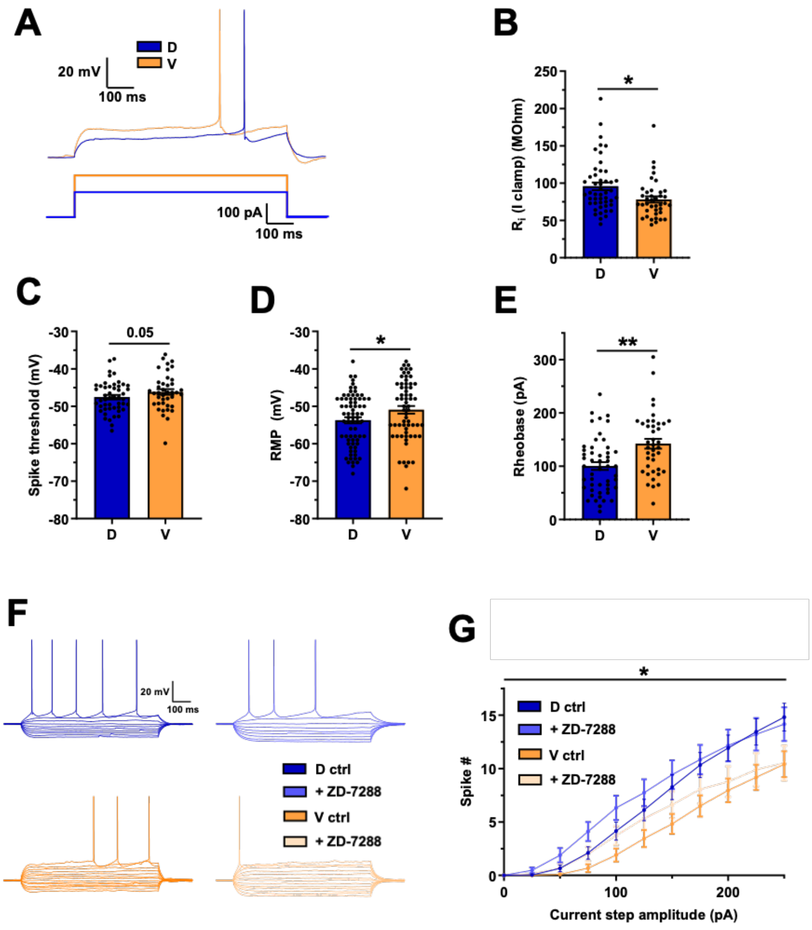


Figure S15 The effect of dorsal-ventral aspect of the hippocampus on intrinsic properties. Data from the rheobase set of experiments. **A)** Voltage deflections in response to a current step of equal size across genotype. Input resistance (3-way ANOVA: aspect main effect: $F_{1, 87}$ = 4.565, P = 0.036) **(B)**, spike threshold (3-way ANOVA: aspect main effect: $F_{1, 89}$ = 3.962, P = 0.05) **(C)**, resting membrane potential (RMP) (3-way ANOVA: aspect main effect: $F_{1, 136}$ = 4.722, P = 0.032) **(D)**, and rheobase (3-way ANOVA: aspect main effect: $F_{1, 90}$= 12.687, P = 0.001) **(E)**, across dorsal-ventral aspects of the hippocampus. 136 cells from 38 animals. **F)** Example dorsal and ventral voltage traces in response to current steps (-150 to 250 pA) before and after application of ZD-7288 20 µM. Spike number across dorsal-ventral aspects of the hippocampus, before and after the application of ZD-7288 20 µM (3-way repeated-measures ANOVA: drug effect: $F_{1, 28}$ = 0.321, P = 0.576. Current step effect: $F_{10, 280}$ = 99.423, P < 0.001. Aspect main effect: $F_{1, 28}$ = 4.890, P = 0.035. Drug x aspect interaction: $F_{1, 28}$ = 0.085, P =

0.773. Drug x step interaction: $F_{10, 280} = 4.126$, $P < 0.001$. Drug x step x aspect interaction: $F_{10, 280} = 1.307$, $P = 0.226$) **(C)**. 37 cells from 18 animals. Summary values depicted as mean ± SEM. * $P < 0.05$, ** $P < 0.01$, *** $P < 0.001$ (3-way ANOVA between subject effect)

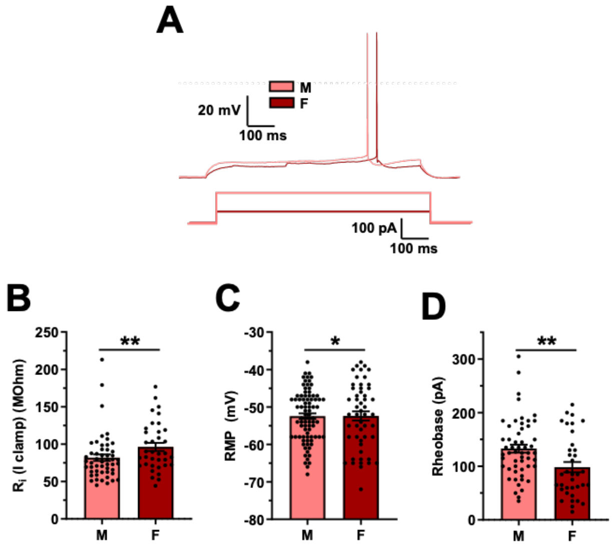


Figure S16 The effect of sex on intrinsic properties. Data from the rheobase set of experiments. **A)** Voltage deflections in response to a current step of equal size across genotype. Input resistance (3-way ANOVA: sex main effect: $F_{1, 87} = 11.739$, $P = 0.001$) **(B)**, resting membrane potential (RMP) (3-way ANOVA: sex main effect: $F_{1, 136} = 5.176$, $P = 0.025$) **(C)**, and rheobase (3-way ANOVA: sex main effect: $F_{1, 90} = 10.581$, $P = 0.002$) **(D)** across dorsal-ventral aspects of the hippocampus. 136 cells from 38 animals. Summary values depicted as mean ± SEM. * P < 0.05, ** P < 0.01, *** P < 0.001 (3-way ANOVA between subject effect)

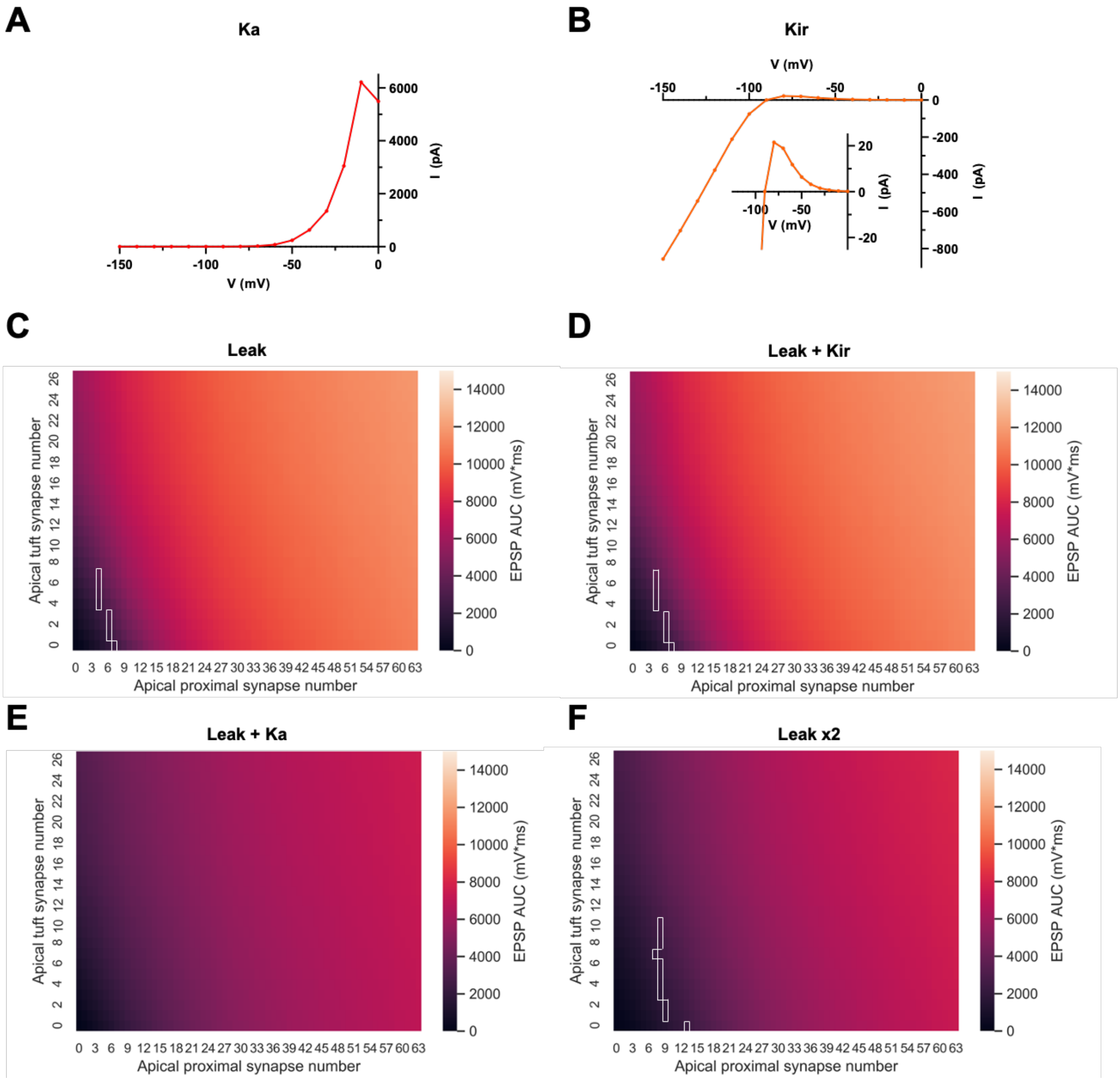


Figure S17 Ka channels abolish and decreased input resistance attenuates dendritic integration irrespective of the number of proximal and tuft dendrites activated. Voltage-current relationships for Ka **(A)** and Kir **(B)** channels, illustrating outward- and inward-rectification, respectively. Inset in B shows the voltage-current relationship for Kir channels at a zoomed in scale for clarity. EPSP area under the curve (AUC) as a function of increasing proximal and tuft synapses from the whole-reconstruction dendritic integration simulations in the presence of leak only **(C)**, leak + Kir **(D)**, leak + Ka **(E)**, and Leak x 2 **(F)**. White boxes indicate the linear-to-supralinear change point.

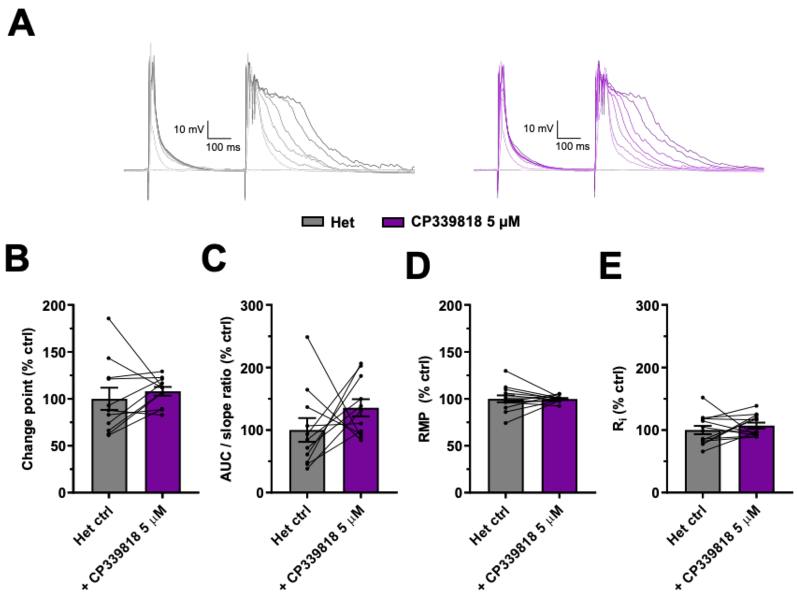


Figure S18 Blocking Kv1.3 and Kv1.4 channels selectively does not affect dendritic integration in the *Dlg2+/-* hets. **A)** Example traces depicting a single EPSP followed by a compound EPSP at increasing stimulation intensities (light to dark) over consecutive recording sweeps before and after CP339818 5 μM. Change point (2-way repeated-measures ANOVA: drug effect: $F_{1, 8} = 2.062$, $P = 0.189$) **(B)**, AUC/slope (2-way repeated-measures ANOVA: drug effect: $F_{1, 8} = 4.553$, $P = 0.065$) **(C)**, resting membrane potential (RMP) (2-way repeated-measures ANOVA: drug effect: $F_{1, 9} = 0.000$, $P = 0.990$) **(D)**, and input resistance (2-way repeated-measures ANOVA: drug effect: $F_{1, 9} = 0.741$, $P = 0.412$) **(E)** as percent of control before and after the after CP339818 5 μM. 13 cells from 8 animals. Summary values depicted as mean ± SEM. * $P < 0.05$, ** $P < 0.01$, *** $P < 0.001$ (3-way ANOVA between subject effect)

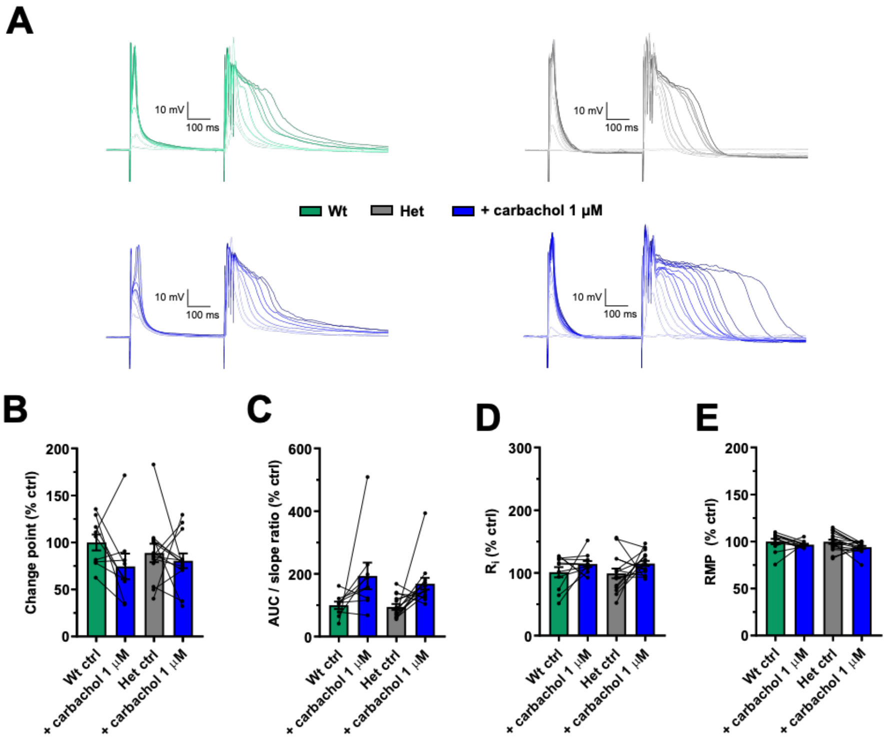


Figure S19 Cholinergic low-dose agonism lowers dendritic integration thresholds in the *Dlg2+/-* hets. **A)** Example traces depicting a single EPSP followed by a compound EPSP at increasing stimulation intensities (light to dark) over consecutive recording sweeps before and after carbachol 1 µM across genotype. Change point (3-way repeated-measures ANOVA: drug effect: $F_{1, 14} = 9.054$, $P = 0.009$. Genotype x drug interaction: $F_{1, 14} = 0.682$, $P = 0.423$) **(B)**, AUC/slope (3-way repeated-measures ANOVA: drug effect: $F_{1, 15} = 28.509$, $P < 0.001$. Genotype x drug interaction: $F_{1, 15} = 0.839$, $P = 0.374$) **(C)**, resting membrane potential (RMP) (3-way repeated-measures ANOVA: drug effect: $F_{1, 21} = 23.469$, $P < 0.001$. Genotype x drug interaction: $F_{1, 21} = 3.254$, $P = 0.086$) **(D)**, and input resistance (3-way repeated-measures ANOVA: drug effect: $F_{1, 21} = 13.203$, $P = 0.002$. Genotype x drug interaction: $F_{1, 21} = 0.001$, $P = 0.974$) **(E)** as percent of control before and after the after carbachol 1 µM across genotype. 32 cells from 15 animals. Summary values depicted as mean ± SEM. * $P < 0.05$, ** $P < 0.01$, *** $P < 0.001$ (3-way ANOVA between subject effect)

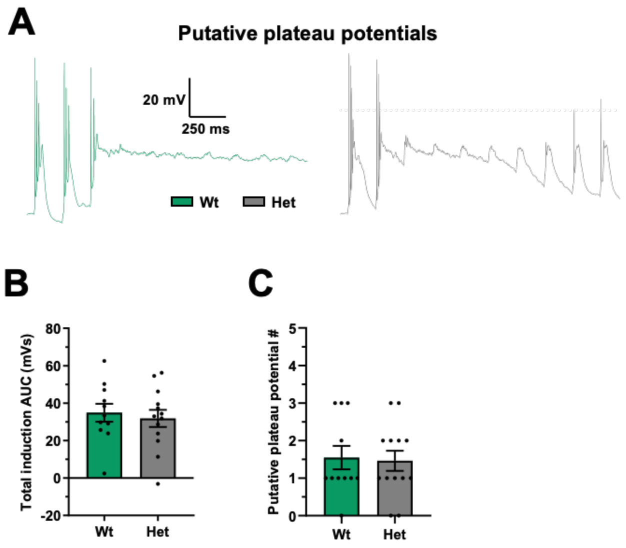


Figure S20 Muscarinic M1 agonism facilitates plateau potential generation in aLTP induction. **A)** example aLTP induction traces showing putative plateau potentials. Total induction area under the curve (AUC) (3-way ANOVA: genotype main effect: $F_{1, 24} = 0.100$, $P = 0.755$) **(B)** and putative plateau potential number (3-way ANOVA: genotype main effect: $F_{1, 24} = 0.002$, $P = 0.963$) **(C)**. Across genotype. 24 cells from 13 animals. Summary values depicted as mean ± SEM. * $P < 0.05$, ** $P < 0.01$, *** $P < 0.001$ (3-way ANOVA between subject effect)
